# Structure of a lasso peptide bound ETB receptor provides insights into the mechanism of GPCR inverse agonism

**DOI:** 10.1101/2023.12.30.573741

**Authors:** Wataru Shihoya, Hiroaki Akasaka, Peter A. Jordan, Anna Lechner, Bethany K. Okada, Gabriella Costa Machado da Cruz, Fumiya K. Sano, Tatsuki Tanaka, Ryo Kawahara, Rajan Chaudhari, Hiroko Masamune, Mark J. Burk, Osamu Nureki

## Abstract

Lasso peptides exhibit a unique lariat-like knotted structure imparting exceptional stability and thus show promise as therapeutic agents that target cell-surface receptors. One such receptor is the human endothelin ET_B_ receptor, which is implicated in challenging cancers with poor immunotherapy responsiveness. The *Streptomyces*-derived lasso peptide, RES-701-3, is a selective inhibitor for ET_B_ and a compelling candidate for therapeutic development. However, meager production from a genetically recalcitrant host has limited further structure-activity relationship studies of this potent inhibitor. Here, we report cryo-electron microscopy structures of ET_B_ receptor in both its apo form and complex with RES-701-3, facilitated by a calcineurin-fusion strategy. Hydrophobic interactions between RES-701-3 and the transmembrane region of the receptor, especially involving two tryptophan residues, play a crucial role in RES-701-3 binding. Furthermore, RES-701-3 prevents conformational changes associated with G-protein coupling, explaining its inverse agonist activity. A comparative analysis with other lasso peptides and their target proteins highlights the potential of lasso peptides as precise drug candidates for G-protein-coupled receptors. This structural insight into RES-701-3 binding to ET_B_ receptor offers valuable information for the development of novel therapeutics targeting this receptor and provides a broader understanding of lasso peptide interactions with human cell-surface receptors.

## Introduction

Lasso peptides are ribosomally synthesized and post-translationally modified peptidic natural products that display a unique lariat-like, threaded and knotted structure^1, 2^ (Extended Data Fig. 1a). The characteristic threaded lasso structure derives from an isopeptide bond connecting the peptide N-terminus to either a glutamic or aspartic acid side chain. Owing to this locked three-dimensional structure, lasso peptides exhibit remarkable stability against heat and proteolytic degradation. The small characterized fraction of the thousands of lasso peptides predicted in bacterial genomes display diverse biological activities, such as enzyme inhibition and receptor blockade leading to antimicrobial anti-cancer, and anti-HIV activities^3^. Lasso peptides appear to occupy a unique functional space, combining the selectivity and potency of larger protein biologics with the low immunogenicity, stability, tissue penetration, and bioavailability of small molecules making them attractive candidates for drug discovery^1, 2^. Despite their great promise, studies of lasso peptides have been hampered by the absence of efficient production systems that enable lasso peptide diversification as well as large scale production. Recent advances in synthetic biology have changed the prospects for lasso peptide drug discovery. In particular, numerous recent studies have demonstrated the heterologous production of lasso peptides in hosts including *Streptomyces*, a well-established bacterial genus for natural product drugs^4^. Furthermore, in 2021, a breakthrough was achieved to successfully produce lasso peptides through a cell-free biosynthesis approach^5^, thus enabling the creation of extensive libraries of these peptides to uncover novel variants with unique characteristics. These advances, alongside structural insights into how lasso peptides target pharmacologically relevant receptors, such as GPCRs, are expected to accelerate the pace of lasso peptide drug discovery.

RES-701 is one of the earliest identified series of naturally occurring bioactive lasso peptides, and includes four variants (RES-701-1 to -4) with almost identical sequences^6^. All four lasso peptide variants function as selective and potent antagonists for the human endothelin ET_B_ receptor^7^, a G-protein coupled receptor (GPCR). ET_B_ constitutes one of the two subtypes of endothelin receptors along with ET_A_ and plays an essential role in vascular regulation^8, 9^. Notably, ET_B_ has been reported to be overexpressed on tumor vascular endothelial cells, leading to immunologically “cold” tumors with attenuated anti-tumor immune responses and resistance to immunotherapy^10, 11^. Consequently, the inhibition of ET_B_ signaling holds promise as a treatment strategy for challenging cancers that exhibit poor responsiveness to existing immuno-oncology agents as a result of ET_B_ overexpression. However, the selectivity and pharmacokinetics of current small molecule ET_B_ antagonists are inadequate^12–14^ (ET_B_/ET_A_ <100x). Hence, inhibitors based on the RES-701 lasso peptides have emerged as highly promising candidates. Nevertheless, our understanding of the structural interactions between lasso peptides and their target molecules is confined primarily to complex structures involving bacterial RNA polymerase bound to antimicrobial lasso peptides^15^, leaving a significant knowledge gap concerning the mechanisms by which lasso peptides target and influence human cell surface receptors. Here, we report the structure of the representative peptide RES-701-3 bound in the pocket of human ET_B_ receptor, shedding light on the mechanisms that govern how the lasso peptide acts on this important GPCR.

## Results

### Production of RES-701-3 and its analogs

Production of lasso peptides by their wild-type bacterial hosts is typically very low (nanograms or micrograms per liter), which is common for secondary metabolites. Thus, RES-701-3 was previously produced by its natural *Streptomyces* strain at 200 micrograms per liter under optimized fermentation conditions on a 1,000 L scale^6^. Such low levels of production are inadequate for drug development and have precluded the advancement of lasso peptides as a therapeutic modality. To gain sufficient quantities for further discovery efforts, a heterologous production host based on *Streptomyces venezeulae* was engineered to produce RES-701-3 and its analogs in ≥ 1 mg/L quantities required to establish structure-activity relationship (SAR) data (Supplementary Table 1, Supplementary Notes). The biosynthetic enzymes for RES-701-3, encoded by the *lasA* (lasso peptide precursor peptide), *lasB2* (peptidase), *lasC* (cyclase), and *lasB1* (RiPP recognition sequence) genes from *Streptomyces auratus* AGR001, were cloned into the pDualP expression vector (Varigen Biosciences) under the control of the NitR promotor (ɛ-caprolactam induction) and conjugated into *Streptomyces venezuelae* ATCC15439 (Extended Data Fig. 1b). Cultivation of this engineered strain in 2 L shaker flasks for 10 days afforded 12 mg/L RES-701-3 (Supplementary Notes). Similarly, single-site analogs of RES-701-3 were produced by introducing the appropriate genetic mutations in the *lasA* gene.

Receptor binding data for RES-701-3 and its analogs were obtained using CHO-K1 cells expressing recombinant human ET_B_ and ET_A_ receptors. Lasso peptides were tested in competition binding assays vs. radiolabeled natural ligand [^125^I]-endothelin-1. Inhibitory constants (Ki) for RES-701-3 and its analogs are shown in Extended Data Table 1. The Ki of RES-701-3 in this assay was 31.5 nM, and comparable to the literature value (4 nM)^6^, while it showed no activity against ET_A_. These data indicate that the RES-701-3 generated in this study has the same biological activity as that produced by its natural *Streptomyces* strain. RES-701-3 showed high selectivity (ET_B_/ET_A_ >1000x), which is superior to the well-known ET_B_ antagonist BQ788 (100x) (Extended Data Table 1).

### Structure determination

Our initial attempts at obtaining diffraction quality crystals for X-ray crystallography of the RES-701-3-bound human ET_B_ receptor were unsuccessful. Thus, we adopted an alternative protein engineering strategy in which the heterodimeric protein calcineurin is fused to a GPCR by three points of attachment at the cytoplasmic ends of TM5, TM6 and TM7^16^. Calcineurin is a calcium- and calmodulin-dependent serine/threonine protein phosphatase composed of the CN-A and CN-B subunits, and its activity is inhibited by the immunosuppressant FK506-FKBP12^17^. For the structural study, we used the thermostabilized ET_B_ receptor used in previous crystallographic studies^14, 18–21^, which contains five thermostabilizing mutations and is truncated after S407^22^. CN-B was inserted into intracellular loop (ICL) 3 of the receptor, and CN-A was fused to its C-terminus via a GS linker (Fig. 1a). This three-point attachment provides a more rigid link with the GPCR transmembrane domain and facilitates particle alignment during data processing, as shown in the structural study of the β_2_AR-CN fusion protein^16^. We successfully purified ET_B_-CN in LMNG/CHS micelles and confirmed its complex formation with FK506 and FKBP12 (Fig. 1b). We performed the cryo-EM structural analysis of the purified ET_B_-CN-FKBP12 complex, and its representative 2D class average visualized all the components of the fusion protein and the FKBP12 (Fig. 1c, d). Eventually, we determined the cryo-EM structures of the apo and RES-701-3-bound ET_B_ receptors at nominal resolutions of 3.3 Å, which allowed us to build a confident model for most of the receptor, CN-A, CN-B, FK506, and FKBP12 (Fig.1e, f, Extended Data Table 2, Extended Data Figs. 2, 3).

**Fig. 1.**
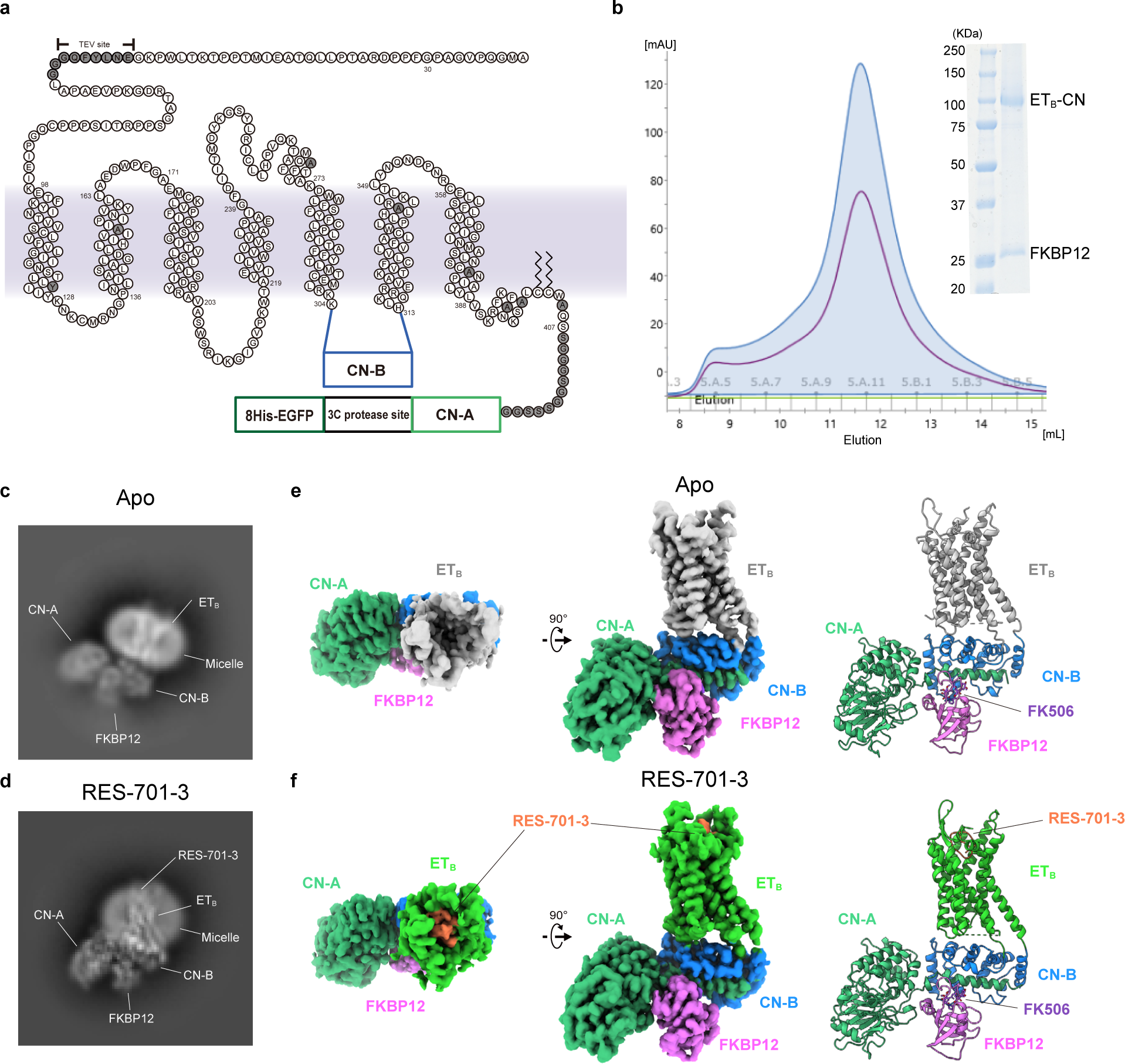
Cryo-EM structure determination of ET_B_ using a three-point fusion strategy. **a**, Concept design of the three-point fusion strategy. **b**, Purification of the ET_B_-CN-FKBP12 complex. **c, d,** Representative 2D cryo-EM averages of the ET_B_-CN-FKBP12 complex in the apo state (**c**) and bound to RES-701-3 (**d**). **e, f,** Cryo-EM density maps and 3D models of the ET_B_-CN-FKBP12 complexes in the apo state (**e**) and complex with RES-701-3 (**f**), viewed from the side and top.

### Architecture of the ETB-CN complex

We first describe the structure of calcineurin and its interaction with the receptor in the apo ET_B_-CN-FKBP12 complex. The structure of calcineurin is essentially similar to the crystal structure of the CN-FK506-FKBP12 complex, while the relative position of CN-A is slightly different (Fig. 2a). Notably, some conformational changes are observed at the junction with the receptor, in which the densities for the linkers between CN-B and receptor are well-resolved (Fig. 2b). In the crystal structure, the area around the C-terminal V169 of CN-B is closed by polar interactions between E18-K83, H13-N89, and R21-V169 (C-terminal carboxylate) (Fig. 2c). In the ET_B_-CN-FKBP12 complex, these interactions are disrupted, and the resulting space is occupied by the ICL3 of ET_B_ fused to the C-terminus of CN-B (Fig. 2b). These local conformational changes in calcineurin allow its receptor coupling.

**Fig. 2.**
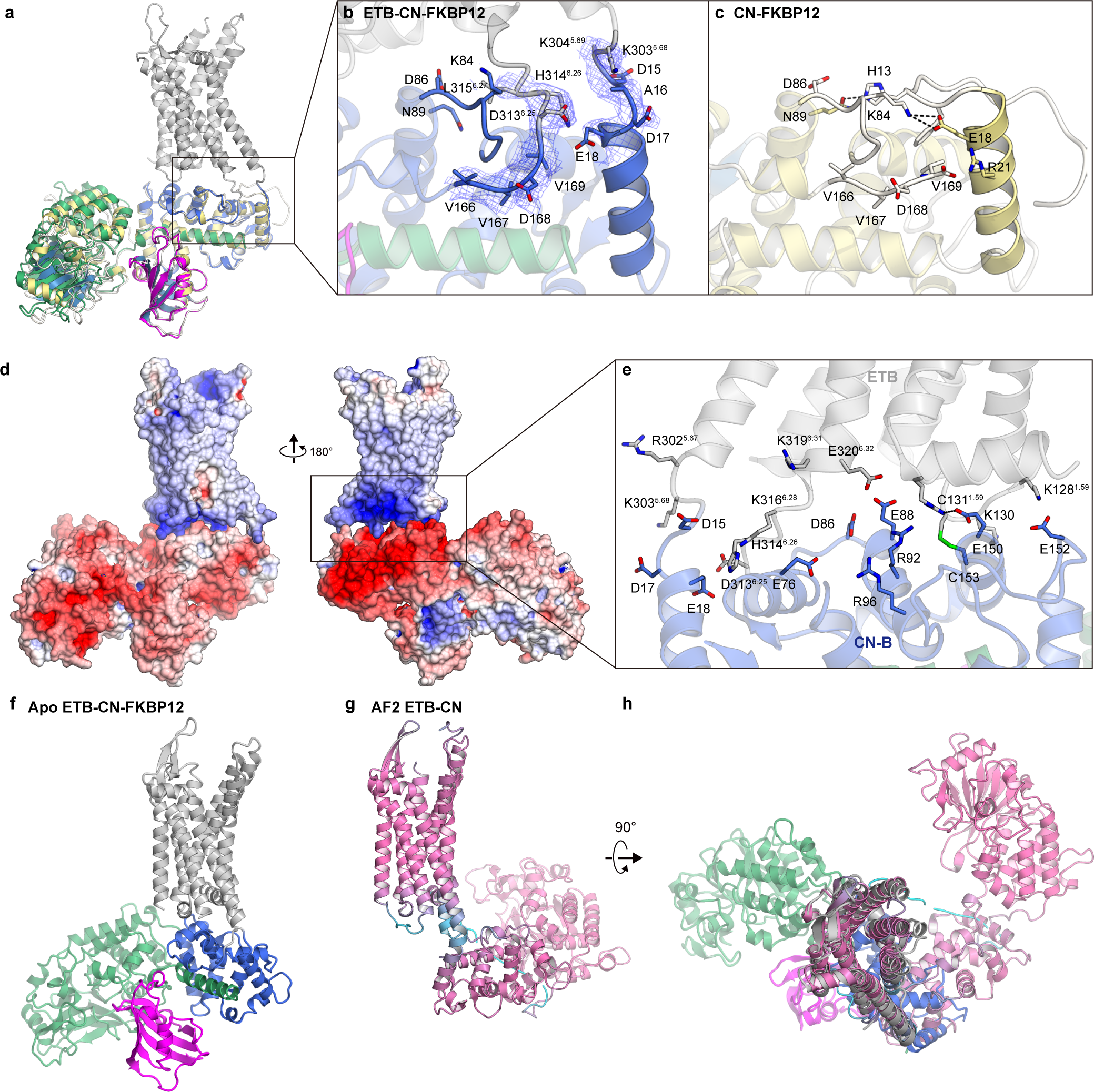
Interactions between ET_B_ and calcineurin. **a,** Structural comparison of the crystal structure of the calcineurin-FKBP12 complex (PDB 1TCO) and the current apo ET_B_-CN-FKBP12 complex. **b, c,** Close-up views of the N-terminus and C-terminus of calcineurin. **d,** Electrostatic surface potentials of ET_B_ and calcineurin in the apo-ET_B_-CN-FKBP12 complex. **e,** Charged residues at the interface of ET_B_ and calcineurin. **f–h**, Structural comparison of the apo-ET_B_-CN-FKBP12 complex and the AF-predicted ET_B_-CN structure. The AF-predicted structure is colored according to PLDDT scores, with higher scores being magenta and lower scores being cyan.

A characteristic feature of the ET_B_-CN-FKBP12 complex is the acquired interaction between ET_B_ and calcineurin. In general, receptors and fusion partners are inherently non-interacting combinations and do not interact outside of the fusion point^23, 24^ (Extended Data Fig. 4a). The intracellular side of ET_B_ is positively charged according to the positive inside rule, whereas CN-B is negatively charged owing to aspartic and glutamic acids exposed on its surface (Fig. 2d). Thus, there are extensive electrostatic interactions between calcineurin and the intracellular face of the receptor (Fig. 2e, Extended Data Fig. 4b). Unexpectedly, C131^ICL^^1^ of ET_B_ is proximal to C153 of CN-B, suggesting a potential intermolecular disulfide bond between them (Fig. 2e). The interaction surface between ET_B_ and calcineurin is 711 Å^2^, which is strikingly larger than that of A2A-BRIL (307 Å^2^) and approaching that of mSMO-PGS (1,010 Å^2^). The mSMO-PGS interface consists primarily of hydrophobic interactions^23^ (Extended Data Fig. 4c), in stark contrast to ET_B_-CN. These interactions stabilize the relative orientations of the receptors and their fusion-partners. The receptor-CN-B interaction is not predicted by AlphaFold^25^, resulting in a large difference in the calcineurin position between the predicted and cryo-EM structures (Fig. 2f–h). This comparison indicates that predicting the structures of GPCR-CN fusions remains challenging.

The receptor structure of the apo ET_B_-CN-FKBP12 complex superimposed well on that of the apo crystal structure of ET_B_-mT4L^18^ (Fig. 3a, b), with a few structural differences. On the intracellular side, the orientations of TM5 and TM6 are different, depending on the fusion partner at ICL3. Moreover, ICL2 is completely disordered due to the steric clash with CN-B (Fig. 3c). On the extracellular side, the β sheet in ECL2 adopts a more open configuration and TM7 moves outwardly by 3 Å (Fig. 3a). Owing to the structural differences in the extracellular regions, the cavity in the apo-ET_B_-CN-FKBP12 complex is wider than that in the crystal structure. The ECL2 conformation is reportedly affected by crystal packing, and thus the cryo-EM structure determined in this study would more accurately reflect the physiological apo state in solution.

**Fig. 3.**
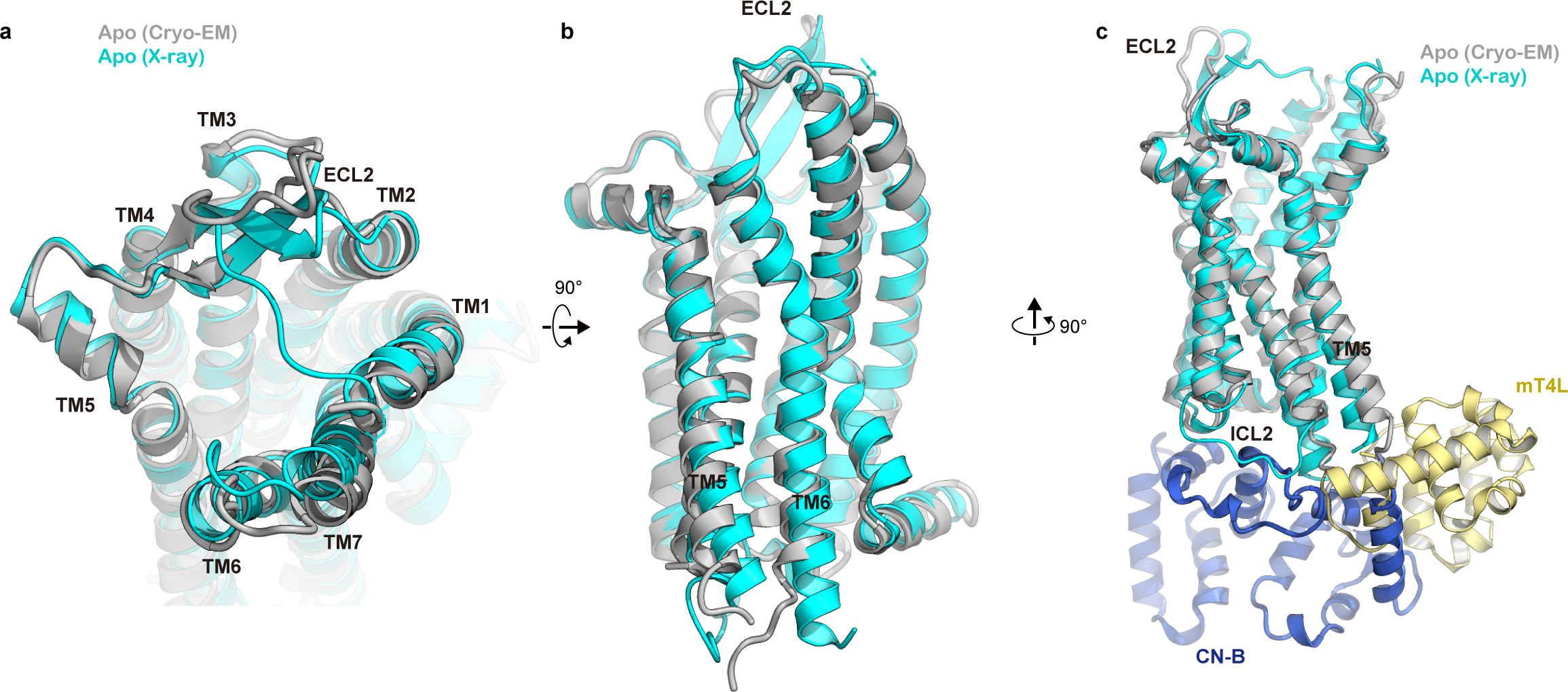
Structural comparison of inactive ET_B_ structures. **a–c**, Superimposition of the apo ET_B_-CN-FKBP12 complex and the apo-crystal structure of ET_B_-mT4L (PDB 5GLI), viewed from the extracellular side (**a**) and the membrane plane (**b and c**). The mT4L and CN-B fusions into ICL3 are shown in (**c**).

### Binding mode of RES-701-3

Within the transmembrane region in the RES-701-3-bound ET_B_-CN-FKBP12 complex, we observed an unambiguous density, enabling us to assign the residues of RES-701-3 except for the C-terminal residue W16 (Fig. 4a, b). RES-701-3 has an isopeptide bond bridging the N-terminal G1 and the carboxyl side chain of D9, forming a nine-residue ring. The C-terminal tail threads through this ring, with sterically locking residues N13 and Y14 on opposite sides of the ring. Between D9 and the locked residue N13, three aromatic residues W10, F11, and F12 create a short hydrophobic loop for binding to the GPCR. Overall, RES-701-3 adopts the typical compact conformation of lasso peptides.

**Fig. 4.**
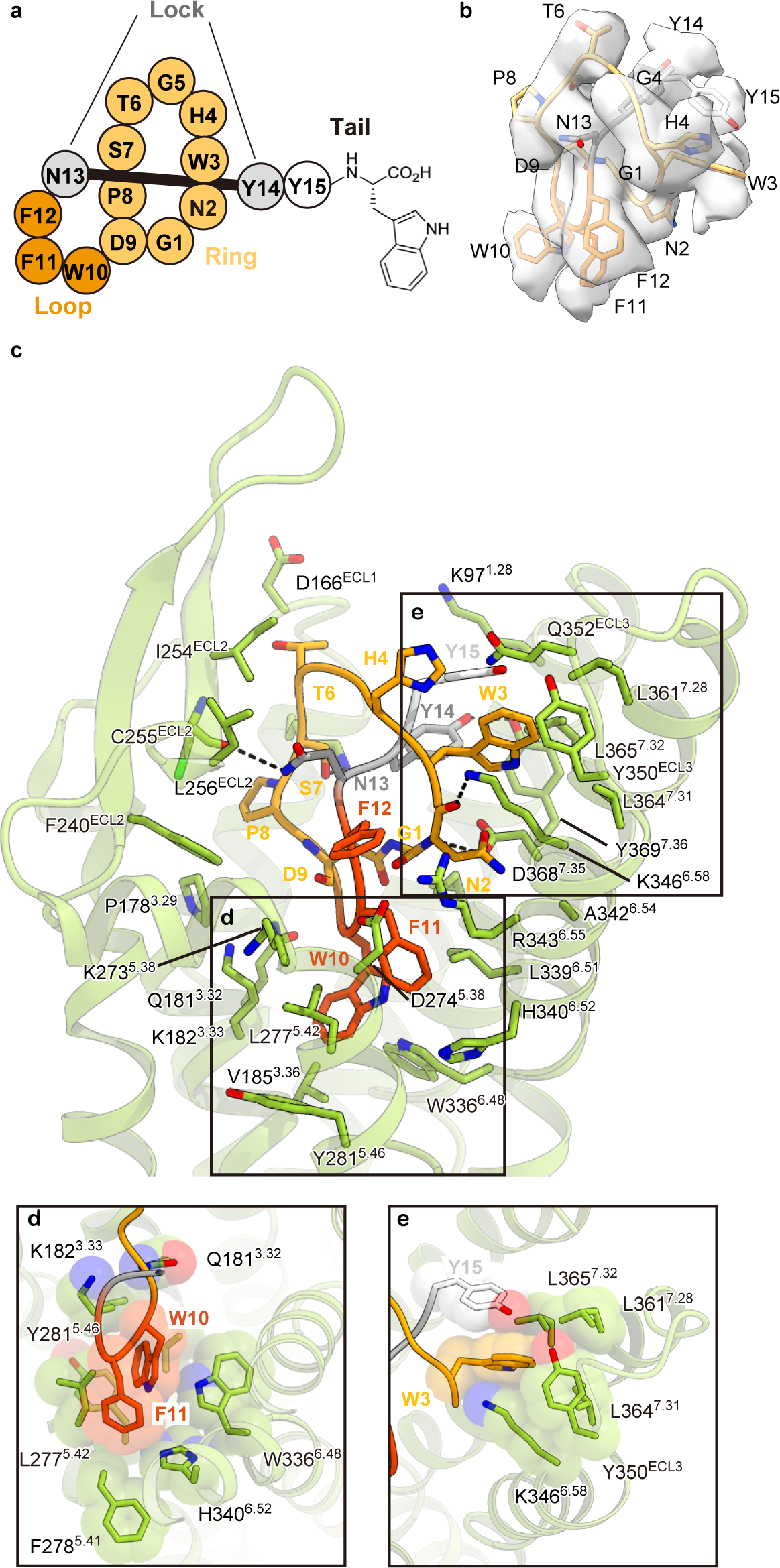
RES-701-3 binding mode. **a,** Schematic illustration of the lasso peptide RES-701-3. **b**, Cryo-EM map of RES-701-3. **c**, Residues involved in RES-701-3 binding within 4.5 Å. Black dashed lines indicate hydrogen bonds. **d, e**, Close-up views of W10 (**d**) and W3 (**e**). Bulky residues are shown as CPK models.

In the complex structure, the loop region of RES-701-3 is oriented toward the transmembrane core, while its C-terminus faces the extracellular milieu. RES-701-3 creates an extensive interaction network with TMs 1–3, TMs 5–7, extracellular loop 1 (ECL) 1 and ECL2 of the receptor (Fig. 4c, Extended Data Table 3, Extended Data Fig. 5a). In total, 31 residues of the receptor interact with RES-701-3 with an interacting surface area of 1,078 Å^2^, accounting for its nM order inhibitory activity and high specificity. W10 and F11 in the loop region form a robust hydrophobic interaction with the inner pocket at the receptor core (Fig. 4d). Moreover, W3 fits into a hydrophobic pocket created by Y15 and bulky residues in TM6 and TM7 (Fig. 4e). Consistently, mutations in aromatic residues W3A, W10A, F11W and Y15A reduced affinity for ET_B_ by more than 100-fold (Supplementary Table 2). H4 and F12, the remaining aromatic residues within RES-701-3, interact to a lesser extent with the receptor, consistent with the relative tolerance for diverse amino acid residue substitutions at these positions. Notably, T6 is in proximity to D166^ECL^^1^, consistent with the 50-fold affinity reduction by the T6E mutation, whereas the mutations of other residues had minimal effects. Overall, the binding mode of RES-701-3 offers comprehensive explanations for prior biochemical findings and our analysis of mutant peptides.

Comparing the apo and RES-701-3-bound ET_B_-CN-FKBP12 complexes, the overall pocket shrinks slightly upon binding, including the inward movements of TMs 1, 3, 7 and ECL2 (Extended Data Fig. 6a). This is also observed when other small molecules bind to ET_B_ (Extended Data Fig. 6b). By contrast, the extracellular portion TM5 is displaced outwardly by 3 Å, a characteristic feature observed only upon RES-701-3 binding. Furthermore, unlike other small molecule inhibitors, RES-701-3 binding does not induce the inward movement of TM6. These structural changes are due to the protrusion of the loop region between TM5 and TM6, which plays an important role in the reception of RES-701-3.

The structure of bound RES-701-3 also provides insights into the activity of previously reported variants of RES-701 isolated from wild type *Streptomyces* cultures. (Extended Data Fig. 5b). As with the C-terminal modifications mentioned earlier, the absence (RES-701-1, -3) versus presence (RES-701-2 and -4) of hydroxylation at W16 only imparts a modest 2-fold impact on receptor binding. This is consistent with the structure where W16 is presented to the extracellular milieu with no specific interactions with the receptor. Similarly, serine 7 (RES-701-3 and -4), compared to alanine 7 (RES-701-1 and -2), imparts a 2-fold improved inhibitory potency. While the structure of RES-701-3 bound ET_B_-CN-FKBP12 suggests a hydrogen bond between serine 7 and K182 of ETB in the current structure this is still quite modest in magnitude.

### Mechanistic insight into inverse agonism

RES-701-3 is not evolutionarily related to the endogenous agonist ligand endothelin-1 (ET-1). Indeed, a structural comparison of the binding modes of ET-1^18, 27^ and RES-701-3 reveals marked differences in the overall binding configurations (Fig. 5a, b). The intramolecular cyclic architecture of ET-1 is mainly recognized in the extracellular region including ECL2, whereas that of RES-701-3 is in the transmembrane region. The C-terminal W21 of ET-1 penetrates into the bottom of the binding pocket, whereas the C-terminal W16 of RES-701-3 is assumed to face the extracellular milieu and is reportedly non-essential for activity. Taken together, the overall binding modes of RES-701-3 and ET-1 are structurally distinct (C-terminus down for ET-1 vs. C-terminus up in RES-701-3). Intriguingly, instead of W16, W10 of RES-701-3 extends to the same depth and position in the binding pocket as W21 of ET-1. Furthermore, the essential W3 interacts with three leucines in TM7, similar to F10 of ET-1. Thus, there is some correspondence between RES-701-3 and ET-1 in terms of the local hydrophobic interactions that are essential for receptor binding.

**Fig. 5.**
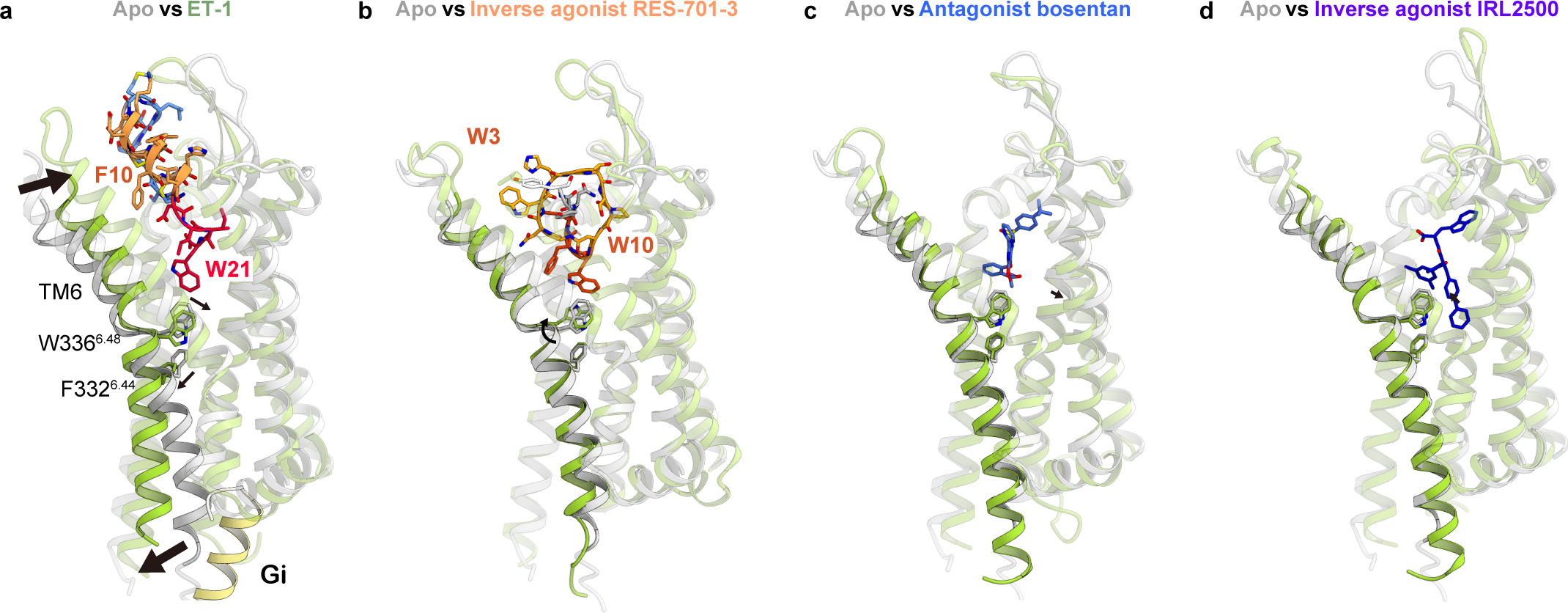
Structural comparison of the ET_B_ receptors. **a–d,** Structural changes upon binding of ET-1 (**a**, PDB 8IY5), RES-701-3 (**b**), bosentan (**c**, PDB 5X93), and IRL2500 (**d**, PDB 6KIQ), focused on TM6. Apo state and compound-bound structures are colored gray and light-green, respectively. Black arrows indicate conformational changes upon drug binding.

Although previous studies have reported that RES-701 lasso peptides function as antagonists for ET_B_, a biochemical analysis using vesicles reconstituted with the purified wild-type ET_B_ showed that they function as inverse agonists^7^. To examine the mechanism of the inverse agonism, we compared the conformational changes upon RES-701-3 binding relative to other drugs (Fig. 5a–d). The binding of the agonist ET-1 induces the inward motions of the extracellular halves of TM6 and TM7, followed by the downward rotation of the W336^6.48^ side chain (Fig. 5a). This rotation of W336^6.48^ induces and propagates the outward rotation of F332^6.44^ within the P^5.50^I/V^3.40^F^6.44^ motif, ultimately resulting in the intracellular opening^27, 28^. The binding of the antagonist bosentan induces only a minor inward movement in TM6^14^ and sterically prevents the rotamer change of W336^6.48^(Fig. 5c). The inverse agonist IRL2500 sandwiches the W336^6.48^ side chain via its aromatic groups, tightly preventing its inward rotation^20^ (Fig. 5d). RES-701-3 occupies the binding pocket more extensively than the small molecule inhibitors, and thereby robustly prevents the conformational change of the receptor. Furthermore, W10 in RES-701-3 rotates the W336^6.48^ side chain outward and away from F332^6.44^ by a direct interaction (Fig. 5b), and thus RES-701-3 binding does not induce the outward rotation of F332^6.44^. Overall, RES-701-3 binding stabilizes the inactive state and prevents the structural transitions of W336^6.48^ and F332^6.44^, plausibly lowering the constitutive activity of the receptor relative to its apo state.

### Insight into ETB selectivity

The RES-701-3 binding site within the transmembrane region is completely conserved between ET_A_ and ET_B_ (Extended Data Fig. 7a). By contrast, the amino acid sequences of ECL1 and ECL2 differ, featuring five inserted residues in ECL1 of ET_A_ (Extended Data Fig. 7a, b). Previous agonist structures indicated that ET_A_ and ET_B_ adopt distinct secondary structures within ECL1 and ECL2, implying their significance in endogenous ligand selectivity^19, 29^. While no antagonist-bound ET_A_ structures have been reported, our findings suggest that RES-701-3 selectively binds to ET_B_ by recognizing the differences in ECL1 and ECL2, in a manner comparable to the endogenous ET_B_-selective ligand ET-3.

## Discussion

In this study, we employed the calcineurin fusion strategy to solve the structure of the RES-701-3-bound ET_B_ receptor. Although this strategy has only been fruitful with β_2_AR, the success described herein with ET_B_ demonstrates that it could be universally applied to structural analyses of GPCRs. Moreover, this fusion strategy allowed the determination of the binding mode of the novel compound RES-701-3, which could not be obtained by X-ray crystallography, showing the utility of the calcineurin fusion strategy for structural determination. RES-701-3 binds differently and has many more points of contact with the receptor, providing an explanation for the very high selectivity of this compound (ET_B_/ET_A_ >1000x) relative to the small molecule BQ-788 (Extended Data Table. 1), which display significantly lower selectivity for ET_B_ receptor (100x). Efficient and highly selective binding to large complex cell surface receptors like GPCRs tends to be challenging for small molecules, underscoring an important advantage for uniquely folded lasso peptides.

Prior to this study, the structures of lasso peptide-target complexes had only been reported for the complex of the antimicrobial peptide MccJ25 with bacterial RNA polymerase^15^ or the siderophore receptor FhuAref^30^. Thus, we compared the binding mode of RES-701-3 with those of the MccJ25-target complexes (Fig. 6a–c), to examine the conserved features of the interactions between lasso peptides and their target proteins. In all the complexes, the lasso peptide becomes entrapped within the binding pocket of the target. In the case of the RNA polymerase-MccJ25 complex, the lasso peptide accesses the secondary channel in a manner typical of substrate binding, with an essential electrostatic interaction between the C-terminal carboxylic acid and a positively charged residue (Fig. 6b). This contrasts with RES-701-3, where the hydrophobic aromatic residues within the loop region play a critical role in the inverse agonist activity. This observation suggests that various segments of the lasso peptide harbor potentials for exerting biological activities. It is noteworthy that comparable cyclic peptides are relatively flat structures that tend to establish superficial interactions with protein surfaces, such as in the crystal structures of thioether-macrocyclic peptides bound to the multidrug and toxic compound extrusion (MATE) transporter^31^(Fig. 6d). The compact structure of lasso peptides renders them exceptionally well-suited for precise targeting of binding pockets, rather than the protein surface. In this context, GPCRs are ideal drug targets for lasso peptides, which offer potential advantages over thioether-macrocyclic peptides. In particular, peptide-activated GPCRs generally have a wider ligand-binding cavity than small molecule-activated GPCRs (e.g., aminergic GPCRs and lipid GPCRs) (Extended Data Fig. 8), making them attractive targets for lasso peptides.

**Fig. 6.**
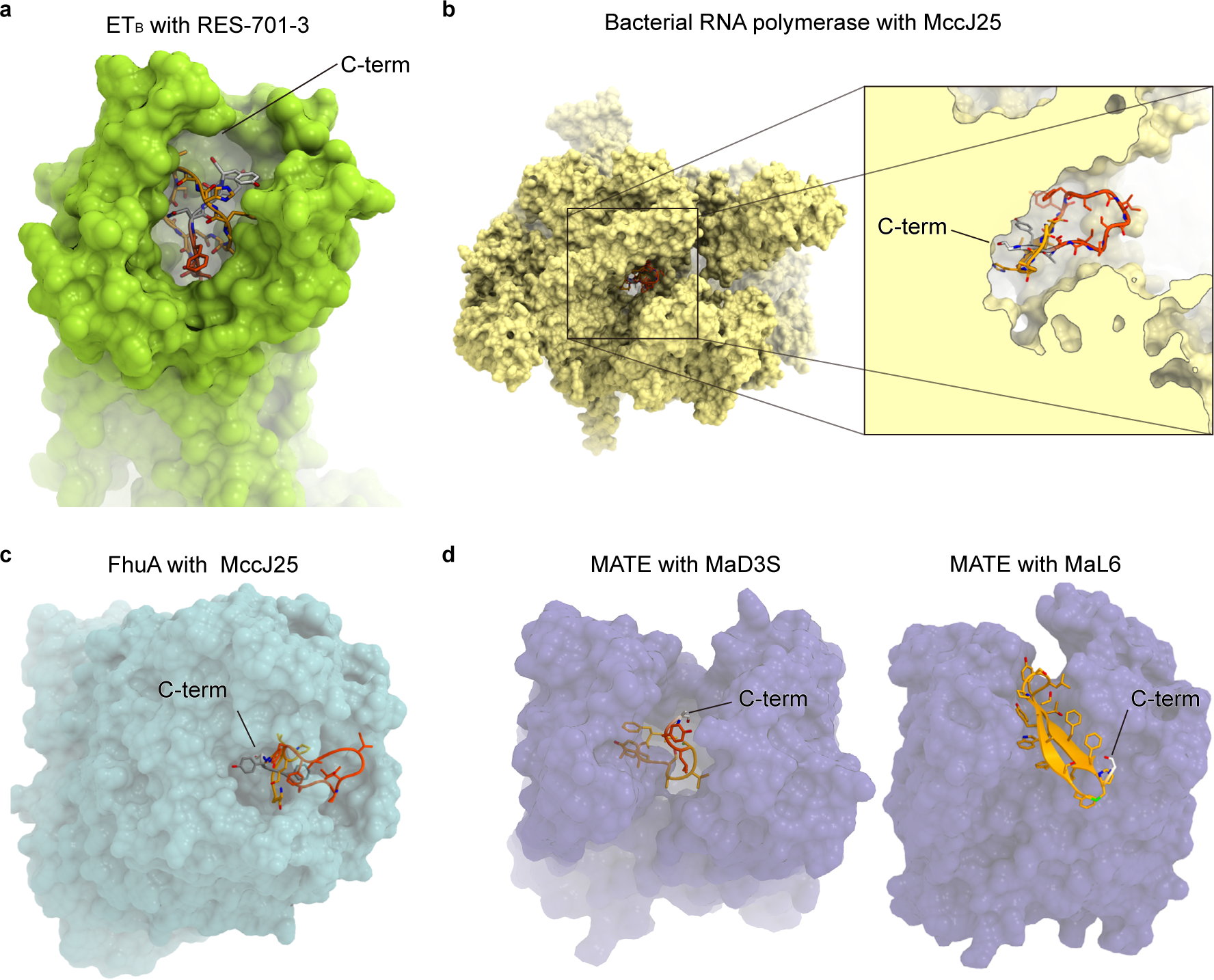
Binding modes of cyclic peptides. **a,** Cavity for RES-701-3. The ET_B_ receptor is shown as a molecular surface. **b**, **c**, Cavities for MccJ25 in bacterial RNA polymerase (PBD 6N60) (**b**) and the siderophore receptor FhuA (PBD 4CU4) (**c**). **d,** Crystal structures of the *P. furiosus* MATE transporter bound to the thioether-macrocyclic peptides MaD3S (PBD 3VVS) and MaL6 (PBD 3WBN).

## Methods

### Competition Binding Experiments

CHO cells were maintained in Kaighn’s F-12K medium supplemented with 10% FB Essence and 2 mM glutamate, under a humidified 5% CO_2_-95% air atmosphere. Cell lines transiently expressing ET_B_ (CHO-ET_B_) were obtained using a mammalian HA-epitope tag expression vector, pHM6 (Roche Applied Science), which carries a cDNA encoding the recombinant human ET_B_ receptor. Each expression vector was introduced into CHO cells by lipofection, using Lipofectamine 2000 (Thermo Fisher, Carlsbad, CA, USA) according to the manufacturer’s instructions. Confirmation of ET_B_ gene expression was confirmed in cell populations by surface staining with antibodies (anti-HA tag AlexaFluor 488 conjugated mouse IgG, R&D Systems, cat # IC6875G) in combination with flow cytometry. Binding experiments were conducted with membranes prepared from the transiently transfected CHO-ET_B_ cells.

CHO-K1 cells expressing recombinant ET_B_ receptors were cultured under standard conditions at 37 °C /5% CO_2_. Cells were collected in ice-cold phosphate buffered saline, pH 7.4 (PBS), and subsequently centrifuged at 500 x g for 5 min at 4 °C. The resulting cell pellet was resuspended in cell lysis buffer containing 5 mM HEPES, pH 7.4, 10 mM EDTA, and 2 mM EGTA, homogenized on ice by Dounce homogenization, and centrifuged (48,000 x g for 15 min at 4 °C). The initial pellet was washed twice by resuspension in 20 mM HEPES, pH 7.4, on ice, and centrifugation (48,000 x g for 15 min at 4 °C). Crude membrane pellets were aliquoted and stored at -80°C prior to use in radioligand binding assays.

The total assay volume in each well of the 96-well microwell plates was 200 µL. Reagent volumes consisted of 3 µL/well of DMSO containing various lasso peptides (with the amino acid sequences described in Extended Data Table 3) prepared at a range of concentrations, 50 µL/well of [^125^I]-endothelin-1 diluted in assay buffer (20 mM HEPES, 10 mM MgCl_2_, 0.2% bovine serum albumin (BSA), pH 7.4), and 150 µl/well of diluted ETAR- or ETBR-expressing membranes prepared in assay buffer. All reagents were combined and incubated for 2 hours at room temperature. Assay incubations were terminated by rapid filtration through Perkin Elmer GF/C filtration plates under vacuum pressure using a 96-well Packard filtration apparatus, followed by washing the filter plates five times with ice-cold assay buffer. Plates were then dried at 45 °C for a minimum of four hours. Finally, 25 µL of BetaScint scintillation cocktail was added to each well and the plates were counted in a Packard TopCount NXT scintillation counter.

Total and non-specific binding were measured in the presence and absence of 10 µM BQ-788. Non-linear regression was used for the analysis of competitive inhibition curves of lasso peptides, and experimentally determined IC_50_ values were used to calculate the dissociation constant (Ki) for each compound, using the Cheng-Prusoff equation.

### Saturation binding studies for determination of radioligand affinity constant (K_d_)

First, 3 µL/well of either DMSO or DMSO containing BQ-788 at a final concentration of 10 µM were added to define total and non-specific binding, respectively. Second, 50 µL/well of assay buffer with serially diluted [^125^I]-endothelin-1 was added. The final concentration of radioligand ranged from 0.015 to 5 nM, calculated based on the stock radioactivity concentration and the specific activity (2200 Ci/mmol). Third, 10 µg/well of diluted membranes were added to initiate the assay. Quadruplicate wells were used for each concentration in the assay. Wells were incubated for 2 hours at room temperature. Assay incubations were terminated by rapid filtration through Perkin Elmer GF/C filtration plates under vacuum pressure using a 96-well Packard filtration apparatus, as described above. The dissociation constant (K_d_) of [^125^I]-endothelin-1 was calculated using non-linear regression analysis of the specific amount of radioactivity bound to the membrane as a function of the radioligand concentration.

### Expression and purification of the ET_B_-CN fusion

The human ET_B_ gene (UniProtKB, Q92633) containing five thermostabilizing mutations^18^ was used as a template. CN-B was inserted in ICL3 between K304 and H313, and CN-A was fused to its C-terminus via a GS linker. The ET_B_-CN fusion was subcloned into a modified pFastBac vector^23^, with an N-terminal haemagglutinin signal peptide and a C-terminal 3C protease recognition site followed by an EGFP-His_8_ tag. The recombinant baculovirus was prepared using the Bac-to-Bac baculovirus expression system (Thermo Fisher Scientific). *Spodoptera frugiperda* Sf9 insect cells (Thermo Fisher Scientific) were infected with the virus at a cell density of 4.0 × 10^6^ cells per milliliter in Sf900 II medium (Gibco), and grown for 48 h at 27 °C. The harvested cells were disrupted by sonication, in buffer containing 20 mM Tris-HCl, pH 8.0, 200 mM NaCl, and 10% glycerol. The crude membrane fraction was collected by ultracentrifugation at 180,000*g* for 1 h. The membrane fraction was solubilized in buffer, containing 20 mM Tris-HCl, pH 8.0, 150 mM NaCl, 1% n-dodecyl-beta-D-maltopyranoside (DDM) (Calbiochem), 0.2% CHS, 10% glycerol, and 2 μM RES-701-3, for 2 h at 4 °C. The supernatant was separated from the insoluble material by ultracentrifugation at 180,000*g* for 30 min, and incubated with TALON resin (Clontech) for 30 min. The resin was washed with ten column volumes of buffer, containing 20 mM Tris-HCl, pH 8.0, 500 mM NaCl, 0.1% lauryl maltose neopentyl glycerol (LMNG) (Anatrace), 0.1% CHS, 0.1 μM RES-701-3, and 15 mM imidazole. After the overnight incubation with the 3C protease (home made), the receptor was concentrated and loaded onto a Superdex200 10/300 Increase size-exclusion column, equilibrated in buffer containing 20 mM Tris-HCl, pH 8.0, 150 mM NaCl, 0.01% LMNG, 0.001% CHS and 0.1 μM RES-701-3. FKBP12 was expressed in *E.coli* and purified by nickel-chromatography, as described previously^17^. The receptor and FKBP12 were mixed at a mol ratio of 1:3. CaCl_2_, FK506, and RES-701-3 were added to achieve final concentrations of 5 mM, 10 uM, and 20 uM, respectively. the receptor was concentrated and loaded onto a Superdex200 10/300 Increase size-exclusion column, equilibrated in buffer containing 20 mM Tris-HCl, pH 8.0, 150 mM NaCl, 0.01% LMNG, 0.001% CHS, 5 mM CaCl_2_, 5 uM FK506, and 0.1 μM RES-701-3. The peak fractions of the ET_B_-CN-FKBP12 complex were collected and concentrated to 12 mg/ml using a centrifugal filter device (Millipore 50 kDa MW cutoff)

### Sample vitrification and cryo-EM single particle analysis

The purified complex was applied onto freshly glow-discharged holey carbon grids (Quantifoil Au 300 mesh R1.2/1.3), which were then immediately plunge-frozen in liquid ethane, using a Vitrobot Mark IV (Thermo Fisher Scientific). Data collections were performed on a 300kV Titan Krios G3i microscope (Thermo Fisher Scientific) equipped with a BioQuantum K3 imaging filter and a K3 direct electron detector (Gatan). Cryo-EM images were collected on a Titan Krios at 300 kV, using a Gatan K3 Summit detector and the EPU software (Thermo Fisher’s single-particle data collection software). Images of the apo state were obtained at an exposure of about 49.983 e^−^ Å^−2^ at the grid, with a defocus range from −0.8 to −1.6 μm. The total exposure time was 2.0 s, with 48 frames recorded per micrograph. A total of 8,547 movies were collected. Images of the RES-701-3-bound state were obtained at an exposure of about 49.236 e^−^ Å^−2^ at the grid, with a defocus range from −0.8 to −1.6 μm. The total exposure time was 2.94 s, with 48 frames recorded per micrograph. A total of 17,005 videos were collected. All acquired movies in super-resolution mode were 2× binned, dose-fractionated, and subjected to beam-induced motion correction implemented in RELION 3.1^32, 33^. The contrast transfer function (CTF) parameters were estimated using patch CTF estimation in cryoSPARC^34^. Particles were initially picked from a small fraction with Gaussian blob picking and subjected to 2D classification. Selected particles were used for training of topaz models^35^. For each full dataset, particles were picked and extracted with a pixel size of 3.32 Å, followed by several rounds of 2D classification to remove ‘junk’ particles. The particles were re-extracted with the pixel size of 1.66 or 1.16 Å and subjected to ab-intio reconstruction and several rounds of hetero refinement in cryoSPARC. Next, two models were obtained by 3D Variability Analysis. These models were used as the initial model, the particles were subjected to several rounds of hetero refinement and non-uniform refinement. For the apo state, the refinement was performed on a further group of particles at an earlier stage, and the 134,931 particles in the best class were reconstructed using non-uniform refinement. For the RES-701-3-bound state, the 95,937 particles in the best class were reconstructed using non-uniform refinement. Those particles were subjected to Bayesian polishing in RELION 3.1^36^, resulting in a 3.33 Å and 3.30 Å resolution reconstruction in the apo state and RES-701-3-bound state, respectively, with the gold-standard Fourier shell correlation (FSC = 0.143). Moreover, the RES-701-3-bound model was refined with a mask on the receptor. As a result, the receptor has a 3.5 Å resolution with a nominal resolution. The overall and receptor focused maps were combined by phenix^37^. The processing strategy is described in Extended Data Fig. 2 and 3.

### Model building and refinement

The quality of the map was sufficient to build a model manually in Coot^38, 39^. The model building was facilitated by the crystal structures of calcineurin (PDB 1TCO)^17^ and apo ET_B_ (PDB 5GLH)^18^. RES-701-3 was manually modeled based on the density. After the model was manually readjusted into the density maps with Coot, it was refined using phenix.real_space_refine (v.1.19)^40^.

## Data Availability

Cryo-EM density maps and structure coordinates have been deposited in the Electron Microscopy Data Bank (EMDB) and the PDB, with the respective accession codes EDM-XXX and PDB YYYY for the apo-state ET_B_-CN-FKBP12 complex, and EMD-XXXX and PDB YYYY for the RES-701-3-bound ET_B_-CN-FKBP12 complex.

## Acknowledgements

We thank K. Ogomori and C. Harada for technical assistance. This work was supported by grants from the Platform for Drug Discovery, Informatics and Structural Life Science by the Ministry of Education, Culture, Sports, Science and Technology (MEXT), and JSPS KAKENHI grants 21H05037 (O.N.), 22K19371 and 22H02751 (W.S.), and 21J20692 (T.T.); ONO Medical Research Foundation (W.S.); The Kao Foundation for Arts and Sciences (W.S.); The Takeda Science Foundation (W.S.); The Uehara Memorial Foundation (W.S.); AMED under Grant Number JP233fa627001 (O.N.); the Platform Project for Supporting Drug Discovery and Life Science Research (Basis for Supporting Innovative Drug Discovery and Life Science Research (BINDS)) from AMED, under grant numbers JP23ama121002 (support number 3272, O.N.) and JP23ama121012 (O.N.).

## Author contribution

W.S. performed the sample preparation and model building. H.A. performed the grid preparation, the cryo-EM data collection, and the single particle analysis, with the assistance by F.K.S. and T.T. R.K purified the FKBP12 protein. P.A.J., A.L., B.K.O., G.C.M.C., R.C., H.M., and M.J.B. performed the production and functional analysis of the lasso peptides. The manuscript was mainly prepared by W.S., H.A., P.A.J., and M.J.B. with assistance from O.N. W.S. and O.N. supervised the project.

