## Extended Data for "Structure of a lasso peptide bound ETB receptor provides insights into the mechanism of GPCR inverse agonism"

**Extended Data Table 1. $ET_A$ and $ET_B$ competition binding data vs [$^{125}$I]-ET-1.**

| Compound | $ET_A$ $IC_{50}$ (nM) | SEM | $ET_B$ $IC_{50}$ (nM) | SEM |
|---|---|---|---|---|
| **RES-701-3** | **>10,000** | **--** | **31.5** | **6.72** |
| **ET-1** | **0.152** | **0.016** | **0.098** | **0.038** |
| **Bosentan** | **6.48** | **1.89** | **116** | **65.0** |
| **BQ-788** | **443** | **98.4** | **3.29** | **0.649** |

$IC_{50}$s are given in nM, with a maximal $IC_{50}$ of 10,000 reported. Standard error of mean (SEM) was reported, with the exception of $ET_A$ data for RES-701-3 which gave values greater than 10,000 each time it was measured.

**Extended Data Table 2. Cryo-EM data collection, refinement and validation statistics.**

| **Data collection** | Apo | RES-701-3-bound |
|---|---|---|
| Microscope | Titan Krios (Thermo Fisher Scientific) | |
| Voltage (keV) | 300 | |
| Electron exposure ($e^-$/Å$^2$) | 49.983 | 49.236 |
| Detector | Gatan K3 summit camera (Gatan) | |
| Magnification | ×105,000 | |
| Defocus range (μm) | -0.8–1.6 | |
| Pixel size (Å/pix) | 0.83 | |
| Number of movies | 8,547 | 17,005 |
| Symmetry | C1 | |
| Picked particles | 5,790,043 | 9,529,132 |
| Final particles | 134,931 | 95,937 |
| Map resolution (Å) | 3.33 | 3.3 |
| FSC threshold | 0.143 | |
| **Model refinement** | | |
| Atoms | 7343 | 7466 |
| **R.m.s. deviations from ideal** | | |
| Bond lengths (Å) | 0.003 | 0.003 |
| Bond angles (°) | 0.568 | 0.642 |
| Validation | | |
| Clashscore | 6.39 | 9.92 |
| Rotamers (%) | 0.37 | 0.00 |
| **Ramachandran plot** | | |
| Favored (%) | 96.77 | 96.16 |
| Allowed (%) | 3.11 | 3.85 |
| Outlier (%) | 0.11 | 0.00 |

**Extended Data Table 3. Interactions of RES-701-3 with ET$_B$.**

| | |
|---|---|
| G1 | D368$^{7.35}$ |
| | I372$^{7.39}$ |
| N2 | A342$^{6.54}$ |
| | R343$^{6.55}$ |
| | K346$^{6.58}$ |
| | D368$^{7.35}$ |
| W3 | K346$^{6.58}$ |
| | Y350$^{ECL3}$ |
| | L361$^{7.28}$ |
| | L364$^{7.31}$ |
| | L365$^{7.32}$ |
| H4 | Q352$^{ECL3}$ |
| G5 | I254$^{ECL2}$ |
| T6 | D166$^{ECL1}$ |
| | W167$^{ECL1}$ |
| | I254$^{ECL2}$ |
| S7 | K161$^{2.46}$ |
| P8 | W167$^{ECL1}$ |
| | P178$^{3.29}$ |
| | F240$^{ECL2}$ |
| | C255$^{ECL2}$ |
| D9 | Q181$^{3.32}$ |
| | I372$^{7.39}$ |
| W10 | Q181$^{3.32}$ |
| | K182$^{3.33}$ |
| | V185$^{3.36}$ |
| | K273$^{5.38}$ |
| | L277$^{5.42}$ |
| | Y281$^{5.46}$ |
| | W336$^{6.48}$ |
| | L339$^{6.51}$ |
| | H340$^{6.52}$ |
| F11 | K273$^{5.38}$ |
| | D274$^{5.39}$ |
| | L277$^{5.42}$ |
| | L278$^{5.41}$ |
| | L339$^{6.51}$ |
| | H340$^{6.52}$ |
| | R343$^{6.55}$ |
| F12 | D274$^{5.39}$ |
| | R343$^{6.55}$ |
| N13 | C255$^{ECL2}$ |
| | L256$^{ECL2}$ |
| Y14 | K97$^{1.28}$ |
| | K161$^{2.46}$ |
| | D368$^{7.35}$ |
| | Y369$^{7.36}$ |
| Y15 | L361$^{7.28}$ |
| | L365$^{7.32}$ |

a

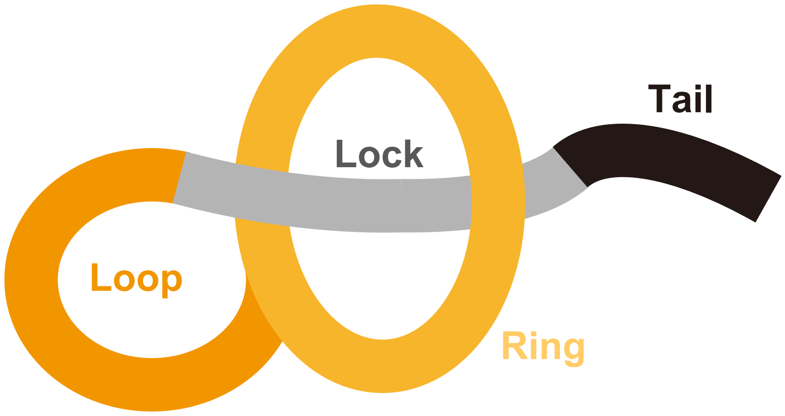


b

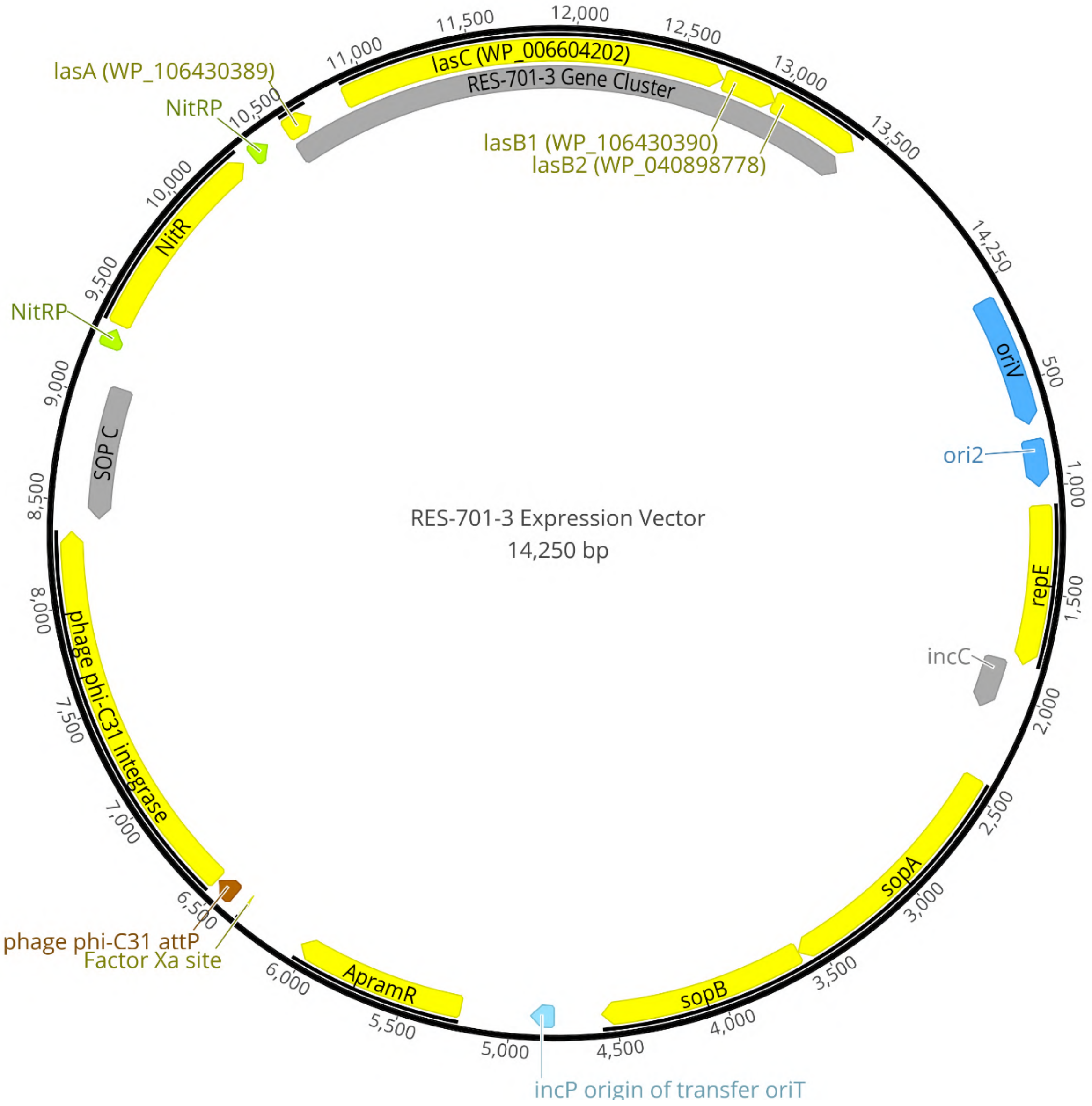


**Extended Data Figure 1| Lasso peptide and vector.**
**a** Schematic diagram of a common lasso peptide. **b** RES-701-3 expression vector. GenBank accession numbers for lasA, lasC las B1 and las B2 are indicated. Abbreviations: **NitRP** – NitR promoter; **oriV** - origin of replication for the bacterial F plasmid; **ori2** - secondary origin of replication for the bacterial F plasmid (also known as oriS); **repE** - replication initiation protein for the bacterial F plasmid; **incC** - incompatibility region of the bacterial F plasmid; **sopA** - partitioning protein for the bacterial F plasmid; **sopB** - partitioning protein for the bacterial F plasmid; **incP origin of transfer oriT** - incP origin of transfer; **ApramR** – apramycin resistance gene; **Factor Xa site** - factor Xa recognition and cleavage site; **phage phi-C31 attp** - attachment site of phage phi-C31; **phage phi-C31 integrase** - integrase from phage phi-C31; **sopC** – sopB binding site.

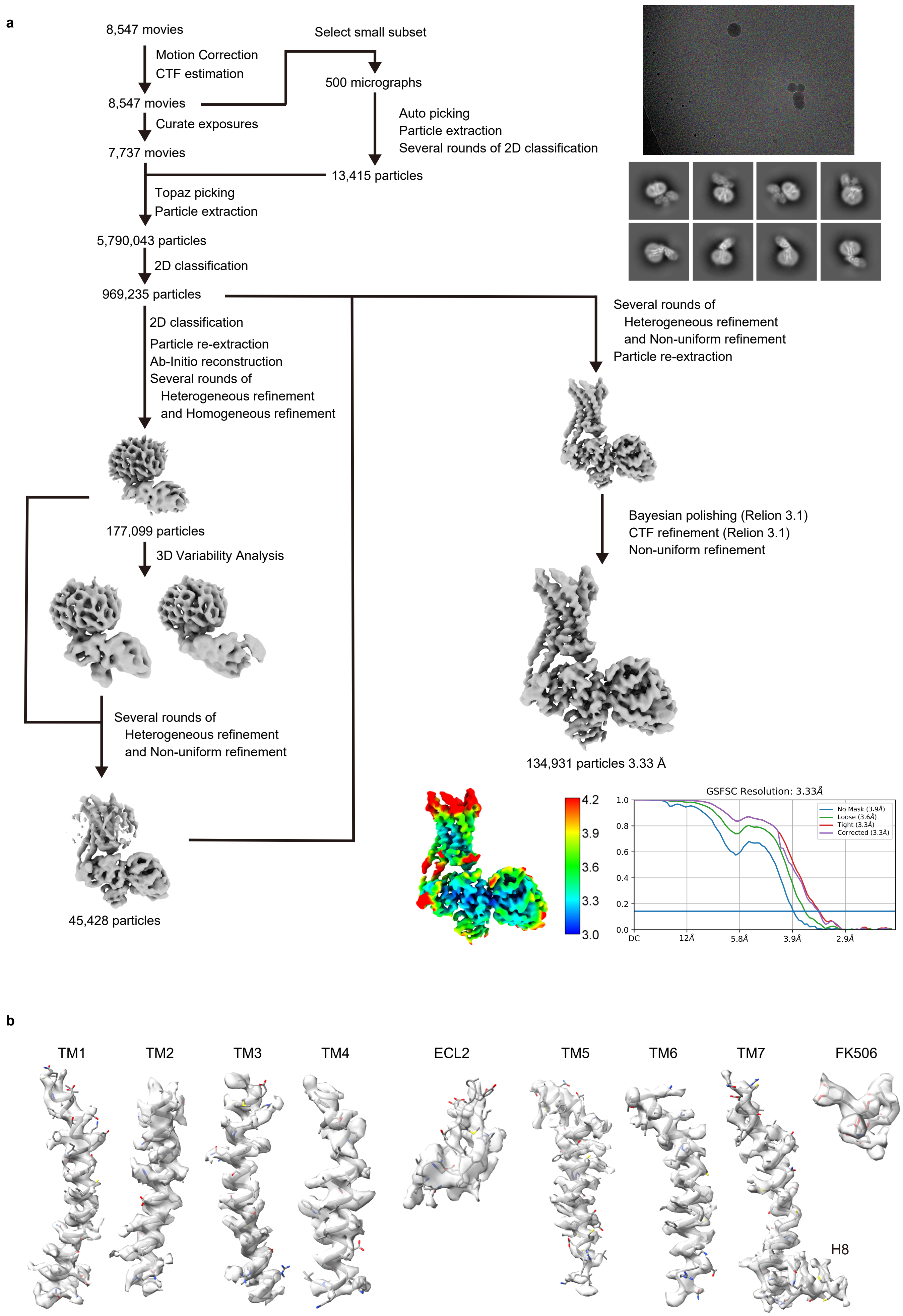


**Extended Data Figure 2| Cryo-EM data processing of apo-state $ET_B$-CN-FKBP12 complex.**
**a** Flow chart of the cryo-EM data processing for apo-state $ET_B$-CN-FKBP12 complex, including particle projection selection, classification, 3D density map reconstruction, local resolution map, and FSC-curves. **b** The cryo-EM density map and model of the receptor are also shown for all seven transmembrane α-helices, ECL2 regions, and FK506. Details are provided in the Methods section.

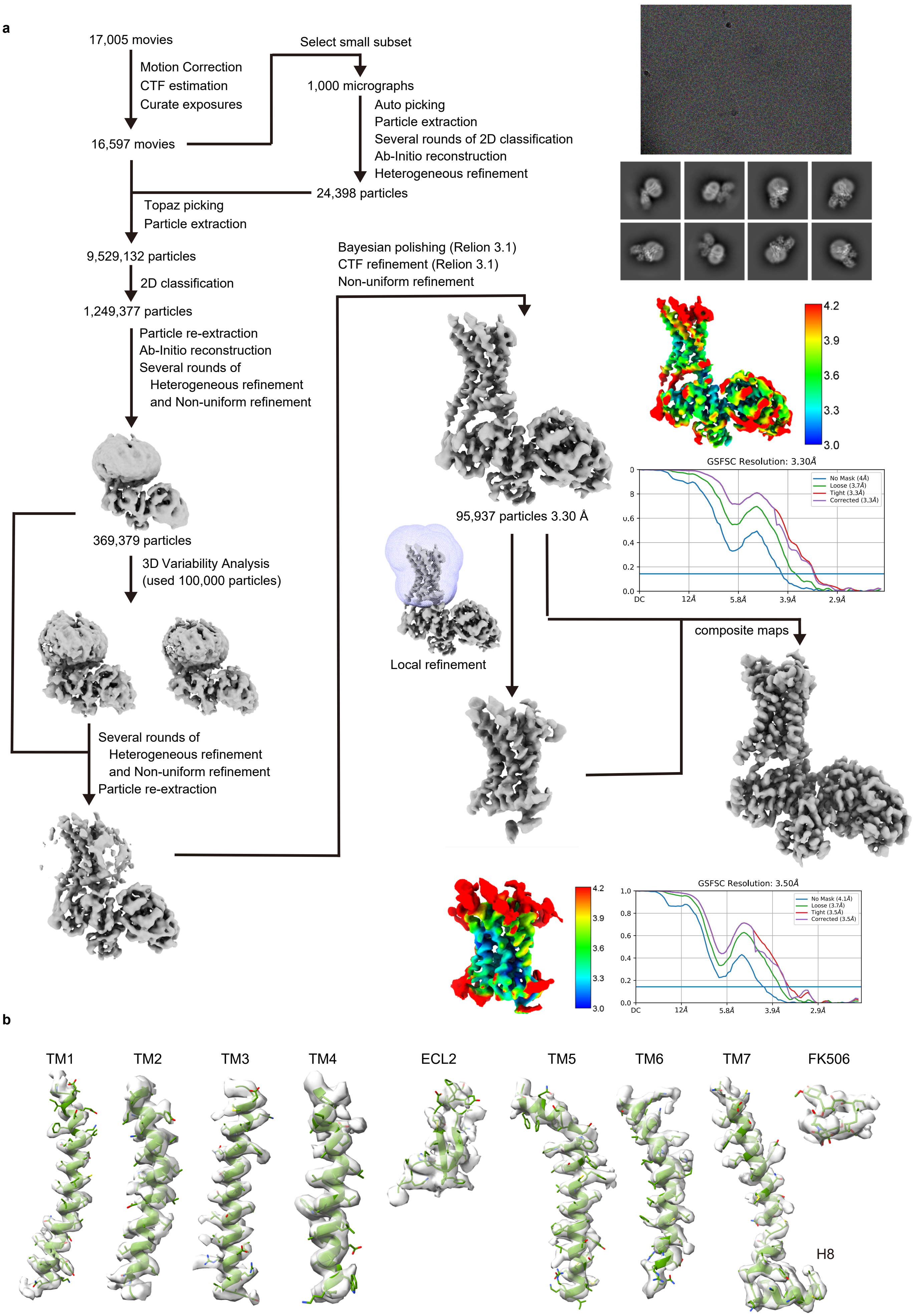


**Extended Data Figure 3| Cryo-EM data processing of RES-701-3-bound $ET_B$-CN-FKBP12 complex.**
**a** Flow chart of the cryo-EM data processing for RES-701-3-bound apo-state $ET_B$-CN-FKBP12 complex, including particle projection selection, classification, 3D density map reconstruction, local resolution map, and FSC-curves. **b** The cryo-EM density map and model of the receptor are also shown for all seven transmembrane α-helices, ECL2 regions, and FK506. Details are provided in the Methods section.

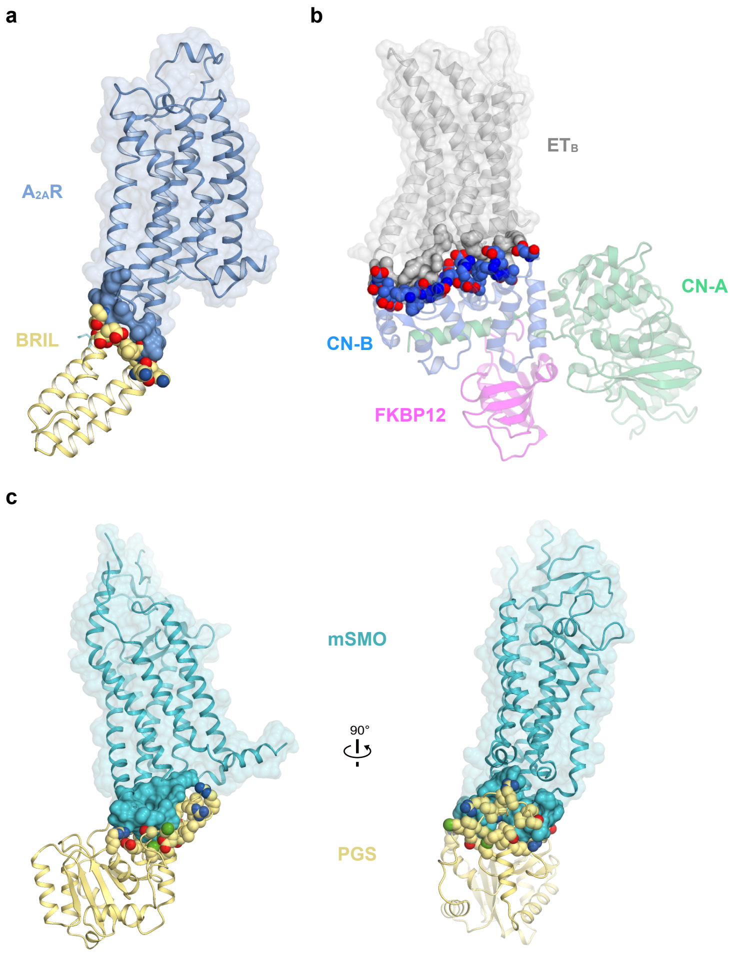


**Extended Data Figure 4 | Comparison of interfaces between GPCRs and their fusion partners.**
**a** $A_{2A}R$ (blue) structure with fused BRIL (yellow) (PDB 7T32). The contact area of $A_{2A}R$ with BRIL is indicated by a blue surface.The residues of BRIL contacting $A_{2A}R$ are indicated by cpk models. **b** 3D models of the $ET_B$–CN-FKBP12 complexes in an apo state. The contact area of $ET_B$ with calcineurin is indicated by a gray surface. The residues of calcineurin contacting $ET_B$ are indicated by cpk models. **c** Two different views of the mSMO-PGS2 structure (PDB 8CXO) showing the relative orientation of the receptor and PGS. The contact area of SMO with PGS is indicated by a blue surface. The residues contacting of PGS with SMO are indicated by cpk models.

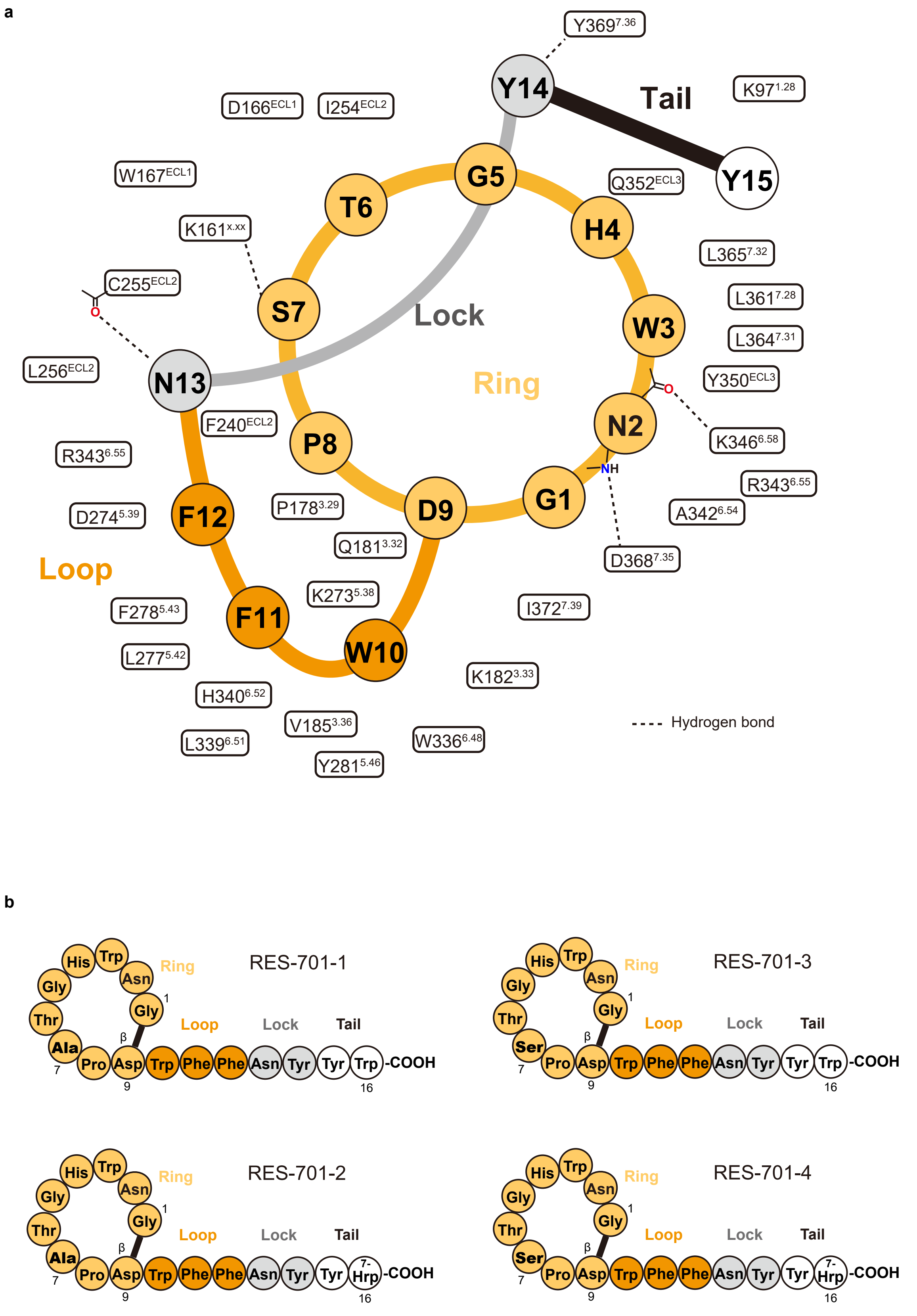


**Extended Data Figure 5 | RES-701-3-receptor interactions.**
**a,** Schematic drawing of the orthosteric pocket. The residues shown here are within a radius of 4.5 Å around RES-701-3. Amino-acid residues of RES-701-3 are represented by capital letters enclosed within circles. Blue and red ovals indicate main chain amide, and carbonyl and carboxyl groups of RES-701-3, respectively. All residues of the $ET_B$ receptor involved in the interactions are indicated by large boxes and amino-acid letters, and the types of interaction are indicated with dotted lines. **b,** Alignment of the amino acid sequences of RES-701s.

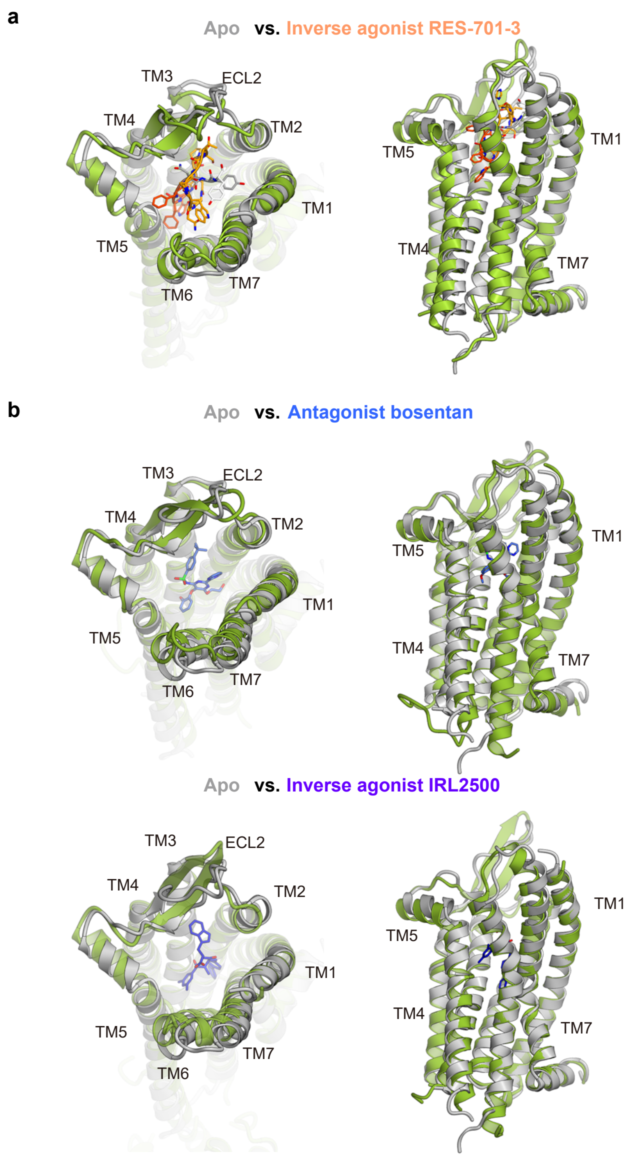


**Extended Data Figure 6 | Structural comparison of inactive $ET_B$ structures.**
**a,** Superimposition of the apo and RES-701-3 bound $ET_B$-CN-FKBP12 complexes.
**b,** Superimposition of the apo $ET_B$-CN-FKBP12 complex and the other inhibitor-bound $ET_B$ structures.

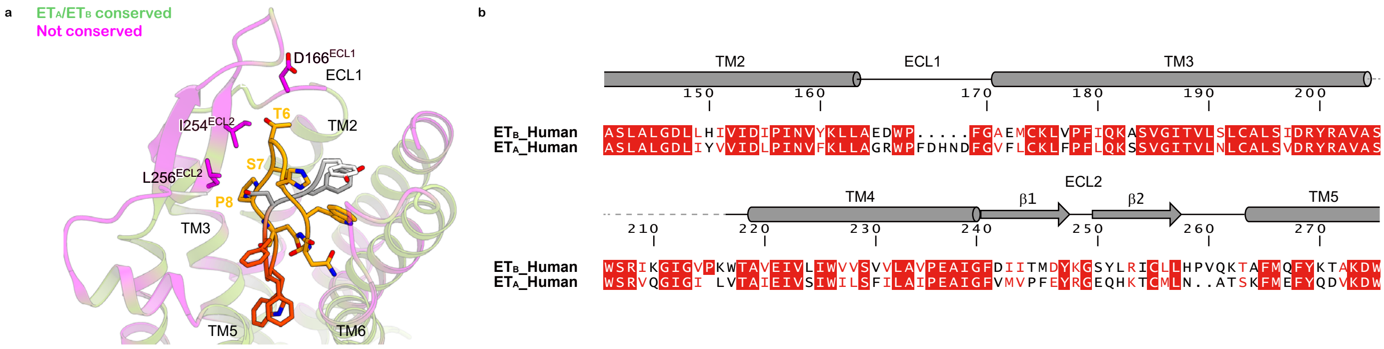


**Extended Data Figure 7 | Conservation of the RES-701-3-binding residues between $ET_B$ and $ET_A$.**
**a** Sequence conservation between the $ET_A$ and $ET_B$ receptors, mapped on the RES-701-3-bound $ET_B$ structure.Conserved and non-conserved residues are colored green and magenta, respectively. Non-conserved residues involved in RES-701-3 binding are indicated by stick models. **b** Alignment of the amino acid sequences of the human $ET_B$ receptor (UniProt ID: P24530) and human $ET_A$ receptor (P25101), focused on ECL1 and ECL2. Secondary structure elements for α-helices and β-strands are indicated by cylinders and arrows, respectively.

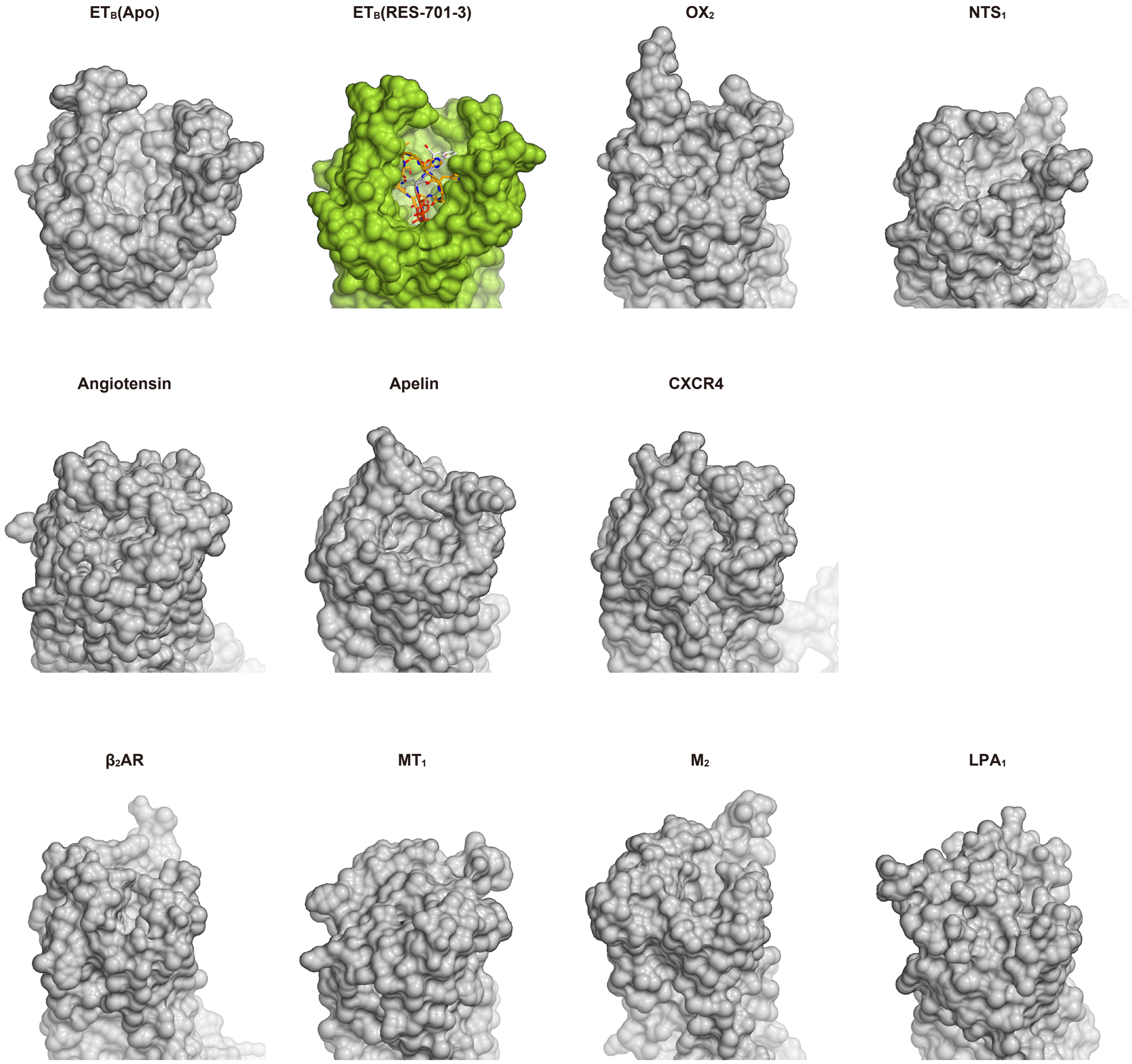


**Extended Data Figure 8 | Binding pockets of novel GPCR structures.**
Molecular surfaces of GPCR structures, viewed from the extracellular side. As shown in the figure, peptide-activated GPCRs: $OX_2$ (4S0V), $NTS_1$ (6ZIN), Angiotensin receptor (4YAY), Apelin receptor (6KNM), and CXCR4 (3ODU) have large binding pockets, compared with those of small-molecule activated GPCRs: $\beta_2AR$ (2RH1), $MT_1$ (6ME2), $M_2$ (3UON), and $LPA_1$ (4Z34), indicating that the former are more suitable targets for lasso peptides.

**Supplementary Table 1**

**Observed m/z and retention times for all RES-701-3 mutants used in competition binding studies.**

| Compound | m/z [M+2H]/2 | Observed m/z | Retention Time* |
|---|---|---|---|
| **RES-701-3** | 1029.932098 | 1030.0 | 7.07 |
| **N2S** | 1016.426648 | 1017.0 | 7.13 |
| **W3A** | 972.41 | 972.6 | 6.79 |
| **H4I** | 1017.944674 | 1018.1 | 8.96 |
| **H4L** | 1017.944674 | 1018.1 | 8.96 |
| **H4M** | 1026.922884 | 1027.5 | 8.77 |
| **H4Y** | 1042.934306 | 1043.5 | 8.48 |
| **H4Q** | 1025.431931 | 1025.6 | 8.09 |
| **H4W** | 1054.44 | 1054.6 | 8.89 |
| **T6V** | 1028.942465 | 1029.1 | 7.26 |
| **T6A** | 1014.93 | 1015.5 | 7.16 |
| **T6S** | 1022.92 | 1023.0 | 7.13 |
| **T6L** | 1035.95029 | 1036.1 | 7.53 |
| **T6I** | 1035.95 | 1036.5 | 7.50 |
| **T6K** | 1043.46 | 1043.6 | 6.52 |
| **T6E** | 1043.93 | 1044.1 | 7.17 |
| **S7E** | 1050.93738 | 1051.0 | 3.66 |
| **S7N** | 1043.43732 | 1043.6 | 7.08 |
| **W10A** | 972.4109983 | 972.9 | 6.16 |
| **F11Y** | 1037.929555 | 1038.1 | 6.79 |
| **F11W** | 1049.437547 | 1050.0 | 7.00 |
| **F12W** | 1049.437547 | 1049.9 | 7.02 |
| **F12H** | 1024.927347 | 1025.1 | 5.73 |
| **F12Y** | 1037.929555 | 1038.1 | 6.62 |
| **F12L** | 1012.94 | 1013.5 | 6.97 |
| **N13S** | 1016.426648 | 1016.6 | 7.02 |
| **Y15L** | 1004.94 | 1005.1 | 7.40 |
| **W16E** | 1001.413738 | 1001.5 | 6.74 |

*Analytical method described in supplementary notes.

**Supplementary Table 2 $ET_B$ Competition binding data vs [$^{125}$I]- ET-1.**

| Compound | IC50 (nM) | SEM | Log fold change | Ref. $IC_{50}$ (nM, RES-701-3) |
|---|---|---|---|---|
| **RES-701-3** | 31.5 | 6.72 | 0 | |
| **N2S** | 105 | -- | -0.60 | 26.4 |
| **W3A** | >10000 | -- | -2.77 | 16.8 |
| **H4I** | 283 | -- | -1.14 | 20.5 |
| **H4L** | 64.0 | -- | -0.49 | 20.5 |
| **H4M** | 17.7 | -- | 0.06 | 20.5 |
| **H4Y** | 26.9 | -- | -0.12 | 20.5 |
| **H4Q** | 16.8 | -- | 0.09 | 20.5 |
| **H4W** | 92.0 | -- | -0.51 | 28.5 |
| **T6V** | 62.5 | -- | -0.18 | 41.0 |
| **T6A** | 30.1 | -- | -0.02 | 28.5 |
| **T6S** | 66.4 | -- | -0.37 | 28.5 |
| **T6L** | 110 | -- | -0.59 | 28.5 |
| **T6I** | 134 | -- | -0.67 | 28.5 |
| **T6K** | 761 | -- | -1.43 | 28.5 |
| **T6E** | 3220 | -- | -2.05 | 28.5 |
| **S7E** | 2280 | -- | -1.39 | 91.9 |
| **S7N** | 145 | -- | -0.55 | 41.0 |
| **W10A** | >10000 | -- | -2.69 | 20.5 |
| **F11Y** | 21.2 | 4.43 | 0.179 | 31.5 |
| **F11W** | >10000 | -- | -2.69 | 20.5 |
| **F12W** | 9.8 | -- | 0.32 | 20.5 |
| **F12H** | 7.9 | -- | 0.41 | 20.5 |
| **F12Y** | 4.0 | -- | 0.71 | 20.5 |
| **F12L** | 14.1 | -- | 0.08 | 16.8 |
| **N13S** | 620 | -- | -0.83 | 91.9 |
| **Y15L** | 3710 | -- | -2.34 | 16.8 |
| **W16E** | 1290 | -- | -1.69 | 26.4 |

$IC_{50}$s are giving in nM, with a maximal $IC_{50}$ of 10,000 reported. Each batch of mutants were assayed in triplicate alongside WT RES-701-1 (WT $IC_{50}$ values are reported in column 5 "Ref. Ki (nM, RES-701-3)"). Log fold change in $IC_{50}$ ($Log(IC50_{RES\text{-}701\text{-}3}/IC50_{mutant})$) was calculated in reference to the RES-701-3 measured in each batch of compounds to directly compare the effect on inhibition. $IC_{50}$s for RES-701-3 and the F11Y mutant were measured multiple dates and $IC_{50}$ values are reported as averages, over 11 and 6 runs, respectively. For these lassos, standard error of mean (SEM) is reported in column 3. Overall, mutations of RES-701-3 tended to reduce the affinity for $ET_B$, especially W3A, W10A, F11W, and Y15L. By contrast, F11Y increased the affinity, suggesting hydrogen bond formation with $H340^{6.52}$. F12Y, F12H, and F12W also increased the affinities. F12 forms van der Waals interactions with $D274^{5.39}$ and $R343^{6.55}$, and thus F12Y and F12H potentially form polar interactions with these residues. Moreover, F12 faces the solvent and has a space that permits the replacement to tryptophan, which would increase the interactions with the receptor. It should be noted that T6K reduces the affinity for $ET_B$ by about 30-fold, although not as much as T6E (100-fold). This is thought to be a steric hindrance effect due to the substitution with bulkier residues.

### Supplementary notes

#### General Methods

Reagents used for molecular biology experiments were purchased from New England BioLabs (Ipswich, MA), Thermo Fisher Scientific (Waltham, MA), Gold Biotechnology Inc. (St. Louis, MO) or Integrated DNA Technologies (IDT, Coralville, IA). Other chemicals were purchased from Sigma-Aldrich (St. Louis, MO). Solid phase extractions resins were purchased from Itochu Corporation. The non-methylating *Escherichia coli* (*E. coli*) donor ET12567 and ET12567 carrying the self-transmissible pUB307 were used for intergeneric transfer of plasmids from *E. coli* to *Streptomyces venezuelae* ATCC 15439. All molecular biology manipulations were conducted using standard plates, vials, and flasks typically employed when working with biological molecules such as DNA, RNA and proteins. High-resolution LC-MS analyses were performed on an Agilent 6530 Accurate-Mass Q-TOF MS equipped with a dual electrospray ionization source and an Agilent 1260 LC system with diode array detector. MS and UV data were analyzed with Agilent MassHunter Qualitative Analysis version 10.0. Preparative HPLC was carried out using an Agilent 1100 purification system (ChemStation software, Agilent) equipped with an autosampler, multiple wavelength detector, a Prep-LC fraction collector and Phenomenex Luna 5µm C18(2) 150x30 mm preparative column. NMR data are acquired using a 600 MHz Bruker Avance III spectrometer with a 1.7 mm cryoprobe. All signals are reported in ppm with the internal DMSO-d6 signal at 2.50 ppm ($^1$H-NMR) or 39.52 ppm ($^{13}$C-NMR). 1D data is reported as s = singlet, d = doublet, t = triplet, q=quadruplet, m = multiplet or unresolved, br = broad signal, coupling constant(s) in Hz[1–8].

**Culture media**

*E. coli culturing media*- Luria-Bertani (LB) liquid [10 g/L casein peptone, 5 g/L yeast extract, 10g/L NaCl, pH 7.0] and solid [10 g/L casein peptone, 5 g/L yeast extract, 10g/L NaCl, 15 g/L agar, pH 7.0] media were used.

*Streptomyces culturing media* - For germination of *Streptomyces* spore stocks, 2 x YT medium [16 g/L tryptone, 10 g/L yeast extract, 5 g/L NaCl] was used. For conjugation and selection in *Streptomyces* strains, mannitol soya flour (MS) solid medium [20 g/L mannitol, 20 g/L soya flour, 20 g/L agar] was used. For culturing of *Streptomyces* strains, tryptone soya broth (TSB) medium [17 g/L pancreatic casein peptone, 2.5 g/L $K_2HPO_4$, 2.5 g/L glucose, 5 g/L NaCl, 3 g/L papain digested soya peptone] and R5 medium [103 g/L sucrose, 0.25 g/L $K_2SO_4$, 10.12 g/L $MgCl_2.6H_2O$, 10 g/L glucose, 0.1 g/L Difco casaminoacids, 2 mL/L trace element solution, 5 g/L Difco yeast extract, 5.73 g/L TES buffer.  For trace element solution, 40 mg/L $ZnCl_2$, 200 mg/L $FeCl_3.6H_2O$, 10 mg/L $CuCl_2.2H_2O$, 10 mg/L $MnCl_2.4H_2O$, 10 mg/L $Na_2B_4O_7.10H_2O$, 10 mg/L $(NH_4)_6Mo_7O_{24}.4H_2O$; after autoclaving, 10 mL/L $KH_2PO_4$ (0.5% w/v), 4 mL/L $CaCl_2.2H_2O$, 15 mL/L L-proline (20% w/v), 7 mL/L NaOH (1N) were used.

**Preparation of electrocompetent ET12567 cells** - a single colony from a freshly streaked plate was used to inoculate a 10 mL LB culture with the antibiotic chloramphenicol (35 µg/mL) in a 50 mL Falcon tube and shaken at 200 rpm at 37 °C. After 16 hours, 500 µL of the overnight culture was used to inoculate a 50 mL LB with chloramphenicol culture in a 250 mL baffled flask, which was shaken at 200 rpm and 37 °C until $OD_{600}$ 0.4. The cultures were transferred to Falcon tubes on ice and centrifuged for 15 minutes at 4 °C. The supernatant was discarded, and the pellet was washed with 10% glycerol. The wash step was repeated twice and after the last one, the pellet was resuspended in 250 µL. The cells were aliquoted in Eppendorf tubes, flash frozen and stored at -80 °C.

**Cloning of RES-701-3 mutants in *Streptomyces venezuelae* ATCC15439.**

The oriT carrying vector plasmid from Varigen Biosciences and gBlock fragments were PCR amplified with Q5® High-Fidelity DNA polymerase using primers with the appropriate homologous sequences between neighboring fragments. Vector fragments were DpnI treated and all PCR fragments were purified with the Zymo DNA Clean and Concentrator® -5 kit. Purified fragments were diluted to 30 fmols and Gibson assembled using NEBuilder® HiFi DNA Assembly Master Mix. After incubation at 50 °C for one hour, 1 µL of the Gibson reaction was electroporated into 50 µL of electrocompetent ET12567 cells.

**RES-701-3 -** The entire genomic region consisting of *lasA*, *lasC*, *lasB1* and *lasB2* genes (Genbank IDs: WP_106430389.1; WP_006604202.1; WP_106430390.1; WP_040898778.1) from *Streptomyces auratus* AGR001 was PCR amplified and cloned into the proprietary pDualP expression vector at Varigen Biosciences (See plasmid map, Extended Data Figure 1), placing the operon expression under the control of the NitR promotor (ϵ-caprolactam induction).

**RES-701-3 mutants** – For all RES-701-3 mutants, a 300 base pair (bp) synthetic gene fragment (gBlock, IDT) encompassing a region from 138 bp upstream of the mutated *lasA* (see Table 3 for mutated nucleotide sequence of *lasA*) to 27 bp downstream, was PCR amplified and Gibson assembled with the corresponding fragments of the *lasACB1B2* pDualP plasmid described above.

**Conjugation -** ET12567 competent cells were transformed (electroporation) with the lasACB1B2 containing pDualP plasmid and plated in LB agar plates with chloramphenicol (35 µg/mL) and apramycin (50 µg/mL) to select for the incoming plasmid. A single colony was used to inoculate 10 mL LB containing chloramphenicol (35 µg/mL) and apramycin (50 µg/mL) and grown overnight at 37 °C and 200 rpm. A single colony of ET12567/pUB307 was also grown in LB plus chloramphenicol (35 µg/mL) and kanamycin (50 µg/mL) for the triparental mating procedure. The overnight cultures were diluted 1:100 in fresh LB plus selective antibiotics and grown at 37 °C until an $OD_{600}$ of 0.4-0.6. The cells were centrifuged at 4000 g washed twice with equal volumes of LB and resuspended in 0.1 volume of LB. On the day of conjugation, 15 µL spores (approximately $10^9$ colony forming units (CFU)) of *S. venezuelae* host strain ATCC15439 were mixed with 100 µL of 2 x YT medium, heat shocked at 50 °C for 10 minutes and allowed to cool. For each conjugation, 100 µL of the heat shocked spores of the host were mixed with 100 µL of both resuspended *E. coli* strains. The mixture was plated out on a MS agar plus 20 mM $MgCl_2$ plate and incubated at 30 °C for 20 hours. After 20 hours, the plates were overlayed with 1 mL of filter sterilized molecular biology grade water containing 1 mg of apramycin and 1 mg of nalidixic acid and distributed evenly with a spreader. After an additional 3 to 4 days, potential exconjugant colonies were picked and restreaked onto a MS plate with apramycin (50 µg/mL) and nalidixic acid (25 µg/mL). After 2 days, colony PCR was used on the restreaked colonies to verify that the incoming plasmid had integrated in the host *Streptomyces* strain.

**Lasso Peptide Fermentation**

**Test production** - Three verified colonies from each conjugation were then used to inoculate a 3 mL TSB culture containing apramycin (50 µg/mL) and nalidixic acid (25 µg/mL) in a 15 mL Falcon tube. After 2 days, 400 µL of the TSB culture was used to inoculate a 10 mL R5 culture with apramycin (50 µg/mL) and ε-caprolactam (0.5% w/v) in a 50 mL bio-reaction tube. The same TSB culture was also used to plate a MS plate with apramycin (50 µg/mL) and nalidixic acid (25 µg/mL). R5 cultures were allowed 7 days to grow before checking for production titers.

**Spore stock preparation** - The 7-day-old MS plate from the colony with the highest titer was used to make a spore stock. This was done by adding 5 mL of filter sterilized molecular biology grade water to the plate and using a sterile cotton swab to displace the spores, suspending them in water. The spore suspension was transferred into a sterile syringe with a wad of cotton wool at the end and filtered into a 15 mL Falcon tube. The spores were centrifuged at 4000 x g for 10 minutes and the supernatant was aspirated with a pipette. The spore pellet was then resuspended in enough 20% glycerol to form an approximately 10% glycerol solution and stored at -80 °C.

**Scale-up production** – Starter cultures were prepared by adding 25 µL of the spore stock to 500 µL of 2 x YT medium and heat shocked at 50 °C for 10 minutes. After cooling down, 500 µL of the mixture was used to inoculate 50 mL of TSB containing apramycin (50 µg/mL) in a 250 mL baffled flask. The culture was shaken at 28 °C at 200 rpm for 1-2 days, then 20 mL of the starter was used to inoculate 500 mL of TSB pre-culture containing apramycin (50 µg/mL) in a 2 L baffled flask. The culture was shaken at 28 °C at 200 rpm for 1 day. 40 mL of the 1-day preculture was used to inoculate 1 L of R5 medium containing apramycin (50 µg/mL) and ε-caprolactam (0.5% w/v). The production culture was shaken at 28 °C at 200 rpm for 10 days.

**Lasso peptide isolation**

**Solid phase extraction of culture broths.**

Lasso peptides were extracted from the whole cell broth by first centrifuging the broth in 750 mL Nalgene bottles (5000 x g). The clarified broth was then added directly to a prepared solid phase extraction (SPE) column as follows:

100g of HP20ss resin (Itochu) was packed into an empty SPE column. The column was washed with 1 L MeOH, then equilibrated with 1 L deionized water. The supernatant extract was loaded onto the SPE resin using a vacuum manifold. The column was washed with 1 L deionized water, then eluted with 1 L 40% MeOH/water, 1 L 50% MeOH/water, 5 x 600 mL 75% MeOH/water, and 1 L 100% MeOH. Each fraction was run on the LC-MS to determine which contained the lasso peptide and approximate the quantity. Fractions containing lasso peptide were pooled and concentrated on a rotary evaporator to remove MeOH, then dried completely on a lyophilizer. Fractions originating from cell pellet material were directly purified on prep HPLC (see below) or were subjected to an additional SPE step.

In some cases, where SPE Fractions were insufficiently clean for HPLC, the fractions were subjected to an addition SPE step as follows:

A 50g HP C18 Redisep column was used for the flash separation. The gradient used DI water as mobile phase A, and HPLC methanol as mobile phase B, with the following program: 0 to 20% B over 5 minutes, 20 to 60% B over 25 minutes, 10 minutes at 60% B, 60 to 75% B over 5 minutes, 10 minutes at 65% B, 75 to 100% B over 2 minutes, and 8 minutes at 100% B. Fractions were run on the LCMS quantification method to determine which contained lasso peptide, and those were pooled and concentrated to dryness on a rotary evaporator and lyophilizer. These fractions were then further purified by preparative HPLC (method below).

**Preparative HPLC** - Preparative HPLC was carried out using an Agilent 1100 purification system (ChemStation software, Agilent) equipped with an autosampler, multiple wavelength detector, Prep-LC fraction collector and Phenomenex Luna 5µm C18(2) 150x30 mm preparative column.  Fractions containing lasso peptides were identified using the LCMS method described above prior to combining and lyophilizing.  Product quality control (QC) was performed on the pooled and concentrated lasso fractions (see Product QC method below).

Preparative HPLC Method:

<u>Column</u>: Phenomenex Luna® preparative column 5 µM, C18(2) 100 Å 150x30 mm

Flow rate: 20 mL/min

Temperature: RT

Mobile Phases: HPLC grade water, MeOH, acetonitrile, isopropyl alcohol, trifluoroacetic acid (TFA), in different percentages and used as gradients

Injection amount: varies

Example method: Solvent A is water with 0.05% TFA, solvent B is acetonitrile with 0.05% TFA. 35.5% B for 20.0 min, then 35.5 to 95% B over 1 minute followed by 95% B for 3 minutes. 5 minute post run equilibration time.

**Sample QC -** The purity of eluted lasso peptide was examined by LC-MS on an Agilent 6460C Triple Quadrupole LC/MS system (LC/TQ) equipped with a Jet stream source (AJS), an Agilent 1290 Infinity II LC system, and a diode array detector (DAD). Where necessary, MSMS fragmentation was used to further characterize lasso peptides based on the rule described in Fouque, K.J.D, et al., ***Analyst***, 2018,**143**, 1157-1170, and to confirm amino acid sequences.

Proton NMR and high-resolution LCMS data were acquired as described above to further confirm peptide structures.

Analytical LCMS Method for purity assessment:

Column: Phenomenex Kinetex 1.7 µm XB-C18 100 A, 50 x 2.1 mm column.

Flow rate: 0.4 mL/min

Temperature: 40 °C

Mobile Phase A: 0.1% formic acid in water (LCMS grade)

Mobile Phase B: acetonitrile (LCMS grade)

Injection amount: 1 µL

HPLC Gradient: 5% B for 1.0 min, then 5 to 50% B over 7 minutes followed by 50 to 95% B over 2 min and 95% B for 2 min. 2 minute post run equilibration time

After a blank subtraction, the peaks from the TIC, UV210, and UV254 signals were integrated, and purity was reported as the Area Sum % for each signal using Agilent MassHunter Qualitative Analysis 10.0.

**High resolution mass spectrometry** - Monoisotopic masses were extrapolated from the lasso peptide charge envelop [$(M+H)^{1+}$, $(M+2H)^{2+}$, $(M+3H)^{3+}$] in the m/z 500–3,200 range using a Agilent 6530 Accurate-Mass Q-TOF MS equipped with a dual electrospray ionization source and an Agilent 1260 LC system using an internal reference (see analytical procedure described above). Both MS and MS/MS analyses were performed in positive-ion mode.

**Nuclear magnetic resonance (mining)** - NMR samples are dissolved in DMSO-d6 (Cambridge Isotope Laboratories). All NMR experiments are run on a 600 MHz Bruker Avance III spectrometer with a 1.7 mm cryoprobe. All signals are reported in ppm with the internal DMSO-d6 signal at 2.50 ppm ($^{1}$H-NMR) or 39.52 ppm ($^{13}$C-NMR).
